# Morphology and ultrastructure of pharyngeal sense organs of *Drosophila* larvae

**DOI:** 10.1101/2025.06.07.657386

**Authors:** Vincent Richter, Tilman Triphan, Albert Cardona, Andreas S. Thum

## Abstract

This study provides a comprehensive ultrastructural analysis of the pharyngeal sensory system in *Drosophila melanogaster* larvae, focusing on the four major pharyngeal sense organs: the ventral pharyngeal sensilla (VPS), dorsal pharyngeal sensilla (DPS), dorsal pharyngeal organ (DPO), and posterior pharyngeal sensilla (PPS). Our analysis revealed 15 sensilla across these organs, comprising four mechanosensory, nine chemosensory, and two dual-function sensilla. We identified 35 Type I neurons (six mechanosensory and 29 chemosensory) and six Type II neurons with putative chemosensory functions. Additional sensory structures, including papilla sensilla and chordotonal organs in the cephalopharyngeal region, were characterized. This detailed mapping and classification of pharyngeal sensory structures completes their structural characterization and provides a foundation for future anatomical and functional studies of sensory perception in insects. This work represents a significant step towards a complete analysis of the larval sensory system, providing new opportunities for investigating how an organism processes sensory information to navigate and interact with its environment.

## Introduction

The larva of the fruit fly (*Drosophila melanogaster*) has emerged as a key model system for investigating how organisms process sensory information (reviewed in Gerber and Stocker 2007; Melcher et al. 2007; Apostolopoulou et al. 2015; Joseph and Carlson 2015; Rimal and Lee 2018; Widmann et al. 2018; Thum and Gerber 2019). Among all sensory modalities, assessing food quality and innocuousness is most critical for survival, particularly in holometabolous insects like *Drosophila*, as its larval stage represents the primary feeding phase during which the organism ingests substantial amounts of food to sustain rapid growth and prepare for metamorphosis (Amrein and Thorne 2005). By taking advantage of light microscopy, volume electron microscopy and 3D imaging analysis, recent studies provided detailed ultrastructural characterization of the various components of the larval *Drosophila* sensory system (Grueber et al. 2002; Python and Stocker 2002; Caldwell et al. 2003; Fishilevich et al. 2005; Kreher et al. 2005; Hwang et al. 2007; Xiang et al. 2010; Kwon et al. 2011; Apostolopoulou et al. 2014; Stewart et al. 2015; Berck et al. 2016; Larderet et al. 2017; Rist and Thum 2017; Miroschnikow et al. 2018; Hartenstein et al. 2019; Hernandez-Nunez et al. 2021; Richter et al. 2024; Schoofs et al. 2024). These descriptions encompass the peripheral sensory system, including the four cephalic sensory organs, the terminal organ (TO), dorsal organ (DO), ventral organ (VO), and labial organ (LO), as well as the enteric nervous system, comprising the esophageal ganglion (EG), hypercerebral ganglion (HCG), and proventricular ganglion (PG). Additionally, they include all external body wall sensilla, proprioceptive chordotonal organs, photoreceptors, and most multidendritic sensory neurons that can be found within the larval body segments. In contrast, a detailed ultrastructural description at the level of individual sensilla and cellular resolution is lacking for the pharyngeal sensory system, despite several studies focusing mainly on the general anatomical organization and the mapping of receptor gene expression (Singh and Singh 1984; Singh 1997; Python and Stocker 2002; Gendre et al. 2004; Colomb et al. 2007; Kwon et al. 2011; Apostolopoulou et al. 2016; Miroschnikow et al. 2018). This gap reflects, at least in part, the technical challenges associated with the three-dimensional visualization of deep internal structures by electron microscopy. These limitations are now increasingly overcome by advances in volume electron microscopy (White et al. 1986; Eberle et al. 2015; Xu et al. 2017; Peddie et al. 2022; Wang et al. 2021; Collinson et al. 2023; Takemura et al. 2023; Winding et al. 2023; Peale et al. 2024). Accordingly, we have conducted an ultrastructural analysis of the four principal sensory organs of the larval pharynx using volume electron microscopy (EM). These organs include the ventral pharyngeal sensilla (VPS), dorsal pharyngeal sensilla (DPS), dorsal pharyngeal organ (DPO), and posterior pharyngeal sensilla (PPS) (Python and Stocker 2002; Gendre et al. 2004).

In larvae, Gendre et al. (2004) comprehensively described the four pharyngeal sensory organs and their developmental fate: the DPS and the DPO develop into the adult labral and ventral cibarial sensory organs (LSO and VCSO), respectively; the posterior PPS give rise to the adult dorsal cibarial sensory organ DCSO; whereas the VPS, derived from the labial segment, undergo apoptosis during metamorphosis. These structures were characterized by detailed transmission electron microscopy and morphological analysis (Stocker and Schorderet 1981; Nayak and Singh 1983, 1985; Singh 1997; Kendroud et al. 2017). However, these earlier studies provided only EM-based schematics and a general organization of pharyngeal sensilla, without the comprehensive high-resolution volumetric analysis now achievable through modern EM methods.

Behavioral, physiological, and functional studies have shown that the larval pharyngeal system is instructing taste-guided behaviors. For instance, a single Gr93a positive neuron pair located in the DPS instructs caffeine-dependent choice behavior and aversive odor-caffeine learning (Apostolopoulou et al. 2016; Choi et al. 2016). A current model, therefore, proposes that pharyngeal sensory neurons are activated upon food ingestion (Apostolopoulou et al. 2015; Miroschnikow et al. 2020; Maier et al. 2021). They instruct initiation or repression of food choice, feeding, and associative food learning through direct contact (Choi et al. 2020). In the case of sugars, sensory input from these neurons is combined with internal metabolic state signals, likely conveyed via the enteric system (Mishra et al. 2013; Hückesfeld et al. 2021; Schoofs et al. 2024).

Furthermore, many of the neurons of the DPS, DPO, and PPS are integrated into the pharyngeal organs of the adult, which contrasts with the general principle that adult sensory structures are born during metamorphosis and suggests that the pharyngeal organs may serve a conserved function throughout development (Gendre et al. 2004). Consistent with this, studies in adult *Drosophila* have demonstrated that the pharyngeal organs detect both chemical and physical properties of ingested food and contribute sensory feedback essential for coordinating the peristaltic motor program during unidirectional swallowing (Yang et al. 2021; Qin et al. 2024). Pharyngeal receptor neurons responsive to sugars facilitate the continuation of feeding bouts (LeDue et al. 2015; Yang et al. 2021), whereas those activated by bitter compounds or elevated salt concentrations robustly suppress food intake (Kim et al. 2017; Chen et al. 2021; Sang et al. 2024). In addition, pharyngeal mechanoreceptors relay information about food texture, enabling dynamic modulation of feeding rates to balance between complete food rejection and excessive consumption (Joseph et al. 2017; Yang et al. 2021; Qin et al. 2024). Together, these mechanisms coordinate nutrient homeostasis and the avoidance of harmful substances. Therefore, a better understanding of the building blocks that make up the pharyngeal sensory system would not only help to understand how insects organize food-guided behavior but also provide additional opportunities for pest control in *Drosophila* and insects in general.

Internal gustatory sensilla located in the cibarium and pharynx have been documented across multiple insect orders, including Diptera, Lepidoptera, Coleoptera, Hemiptera, Hymenoptera, Orthoptera, and Odonata (see Table 1S1; Ludwig 1949; Ranade 1967; Galić 1971; Rice 1973; McIver and Siemicki 1981; Nayak and Singh 1983; Lee and Craig 1983; Kirby et al. 1984; Messchendorp et al. 1998; Glendinning et al. 2002; Gendre et al. 2004; Vegliante 2005; Hückesfeld et al. 2010; Rebora et al. 2014; Baik and Carlson 2020; Ortega-Insaurralde et al. 2024). Across these taxa, a common functional theme emerges: the evaluation of ingested material before and during food uptake, often through organs that combine gustatory and mechanosensory roles (Table S1). In *Drosophila* larvae and adults, the pharyngeal organs therefore likely reflect a conserved internal sensory strategy for assessing food quality, texture, and suitability, even though the number, morphology, and organization of sensilla differ between developmental stages (Gendre et al. 2004). Comparable roles are inferred in *Apis mellifera*, where the epi- and hypopharyngeal sensilla are thought to support nectar and pollen evaluation (Galić 1971; Brito Sanchez 2011), and in lepidopteran larvae, where epipharyngeal or cibarial sensilla likely contribute to food acceptance and host-plant choice (Glendinning et al. 2002; Vegliante 2005). In hemipterans such as *Rhodnius prolixus*, the pharyngeal organ is associated with food recognition in the context of blood feeding, indicating a similar requirement for internal evaluation in a distinct ecological niche (Ortega-Insaurralde et al. 2024). In other dipterans, the internal sensory system is adapted to different feeding ecologies: in *Aedes aegypti*, cibarial sensilla are linked to blood-feeding behavior in adult females (McIver and Siemicki 1981; Lee and Craig 1983; Baik and Carlson 2020), which require a blood meal for egg production, whereas in larval *Musca domestica* and *Calliphora vicina,* mechanosensory roles of the pharyngeal organs were supposed, with limited or no direct evidence for strong gustatory control of feeding (Ludwig 1949; Ranade 1967; Hückesfeld et al. 2010). This comparison suggests that pharyngeal and cibarial organs can be broadly conserved as internal sensors for food evaluation, while their sensory focus shifts with diet, life history, and ecological niche. At the same time, the diversity of terms used for these structures likely reflects both conserved components of the insect oral sensory system and lineage-specific modifications of the mouthpart complex.

**Table 1:**
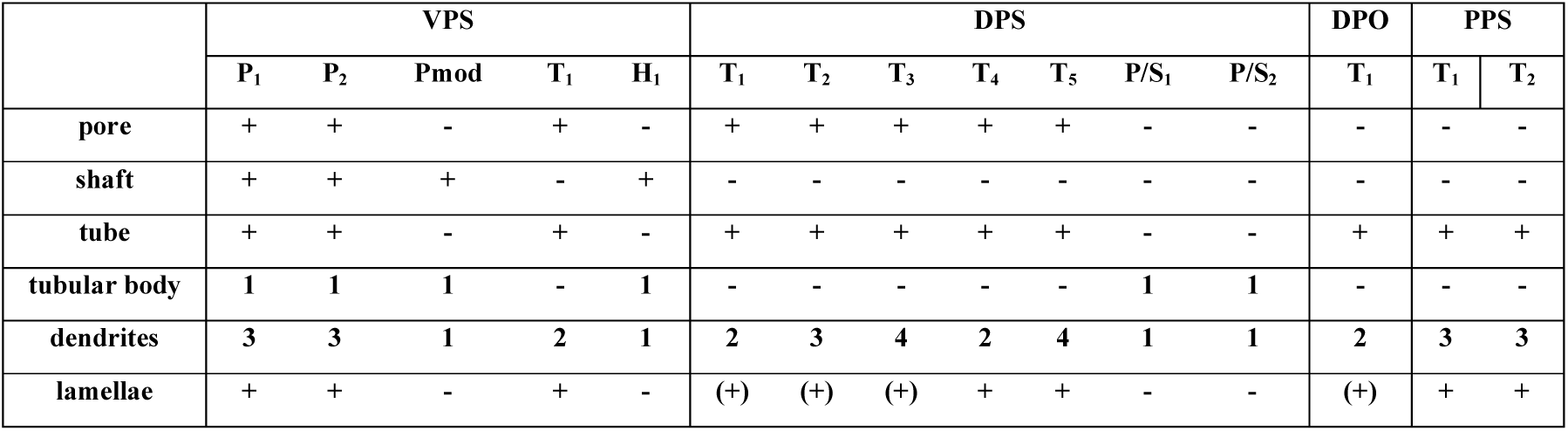
Structural properties of pharyngeal sensilla. Abbreviations: VPS - ventral pharyngeal sensilla; DPS - dorsal pharyngeal sensilla; DPO - dorsal pharyngeal organ; PPS - posterior pharyngeal sensilla; P – papillum sensillum; Pmod – modified papillum sensillum; T – pit sensillum; H – hair sensillum; P/S – papilla/spot sensillum **+** = structure present; (**+)** = structure weakly present; **-** = structure not present.

In this study, we define the morphological types of pharyngeal sensilla based on their ultrastructure, utilizing scanning transmission electron microscopy (STEM) of a larval body volume, which enables the automated acquisition of serial sections at high resolution (Figure 1). The morphological characterization of the pharyngeal sensory structures is based on six criteria: (a) the presence of a terminal pore; (b) the presence of a sensillum shaft; (c) the presence of a cuticle tube; (d) the presence of a tubular body; (e) the number of dendrites; (f) lamellation of the thecogen cell (Table 1). We establish a comprehensive neuron-to-sensillum map for all four pharyngeal organs, identify previously undescribed neurons, and provide a precise classification and unified nomenclature. This work represents a significant step towards a complete analysis of the larval sensory system, providing new opportunities for investigating how *Drosophila* larvae process sensory information to navigate and interact with their environment. Furthermore, the detailed ultrastructural atlas established here provides a foundation for future comparative studies examining the morphological and functional organization of pharyngeal sensory systems in adult *Drosophila* and across other insect species, potentially revealing conserved design principles underlying internal taste sensation and feeding control.

**Figure 1.**
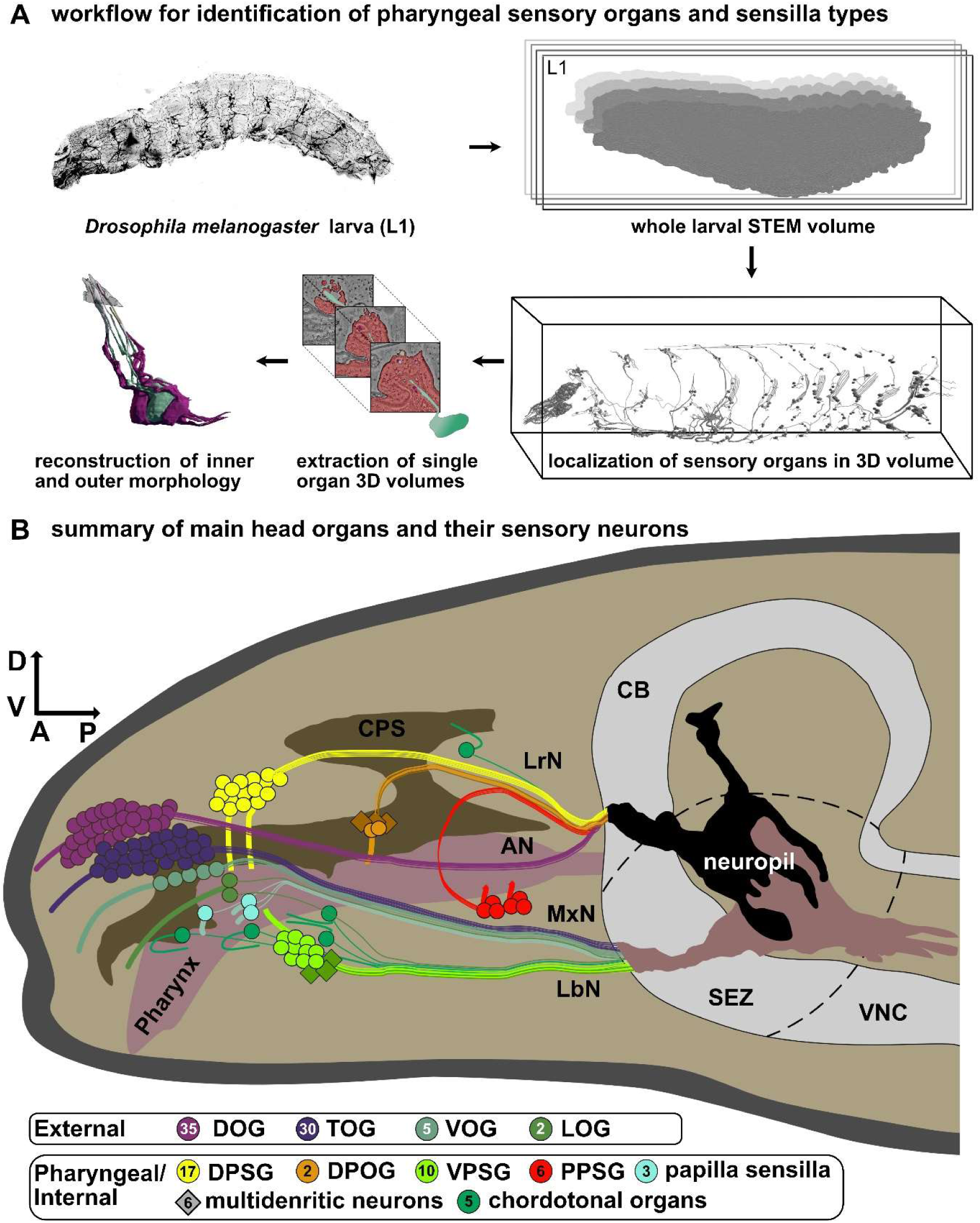
Localization and analysis of the cellular configuration of the larval pharyngeal sensilla. **(A)** Depiction of the workflow for identification of larval pharyngeal sense organs and sensilla types. A previously generated scanning transmission EM (STEM) volume of a whole first instar *Drosophila melanogaster* larva (Peale et al. 2024) was used to localize all cells associated with the larval sensilla using CATMAID (Saalfeld et al. 2009). Smaller volumes were extracted from this volume to generate single organ or sensillum 3D volumes. These volumes were analyzed and manually reconstructed to describe and classify the types of pharyngeal sensilla and their corresponding sensory neurons. **(B)** Schematic drawing of the main external and pharyngeal organs of the larval head region. The projections via the main nerves into the subesophageal zone (SEZ) are visualized. 3D reconstructions of sensilla and sensory neurons were executed from the distal end to the ganglion. Axonal projections were not reconstructed. The numbers in the circles indicate the number of neurons in the respective ganglion. The pharynx (ocher), brain (grey), main external head organs and pharyngeal organs are highlighted. These organs are the dorsal organ (DO; purple), the terminal organ (TO; blue), the ventral organ (VO; light blue), the labial organ (LO; turquoise), the dorsal pharyngeal sensilla (DPS; yellow), the dorsal pharyngeal organ (DPO; orange); ventral pharyngeal sensilla (VPS; bright green) and posterior pharyngeal sensilla (PPS; red). Abbreviations: D - dorsal; V - ventral; A - anterior, P - posterior, L1 – first instar, CB -central brain, SEZ – subesophageal zone, VNC – ventral nerve cord, AN – antennal nerve, MxN – maxillary nerve, LrN – labral nerve, LbN – labial nerve; CPS – cephalopharyngeal skeleton; DOG - dorsal organ ganglion; TOG - terminal organ ganglion; VOG - ventral organ ganglion; LOG - labial organ ganglion; DPSG - dorsal pharyngeal sensilla ganglion; DPOG - the dorsal pharyngeal organ ganglion; VPSG - ventral pharyngeal sensilla ganglion; PPSG - posterior pharyngeal sensilla ganglion

## Results and Discussion

### General

Pharyngeal sensilla share a common basic configuration typical of insect sensilla. They generally contain single or multiple bipolar Type I neurons (Zacharuk and Shields 1991) surrounded by support cells (Schmidt and Berg 1994; Keil 1997; Klowden 2007; Prelic et al. 2021). In contrast to multidendritic Type II neurons, these neurons extend only one dendrite from the cell body toward the outer cuticle or pharyngeal lumen (Klowden 2007). The dendrites comprise inner and outer dendritic segments, separated by the ciliary constriction (Altner and Loftus 1985). The outer segments are immersed in sensillum lymph, enclosed by the dendritic sheath and thecogen (sheath building) cell (Slifer 1970; Altner and Prillinger 1980; Zacharuk 1980; Steinbrecht 1984; Zacharuk and Shields 1991) (e.g., Figure 2B; Figure 2S1-S5 D; Figure 2S5 D; Figure 3S3-S7 C; Figure 4S1-S3 B). Specific structural features indicate sensory function. Mechanosensory Type I neurons typically display a tubular body - a dendritic tip densely packed with microtubules (Thurm 1964; Keil 1997) (e.g., Figure 2B; Figure 2S1-S4 E; Figure 3S1-S2). Chemosensory and especially gustatory sensilla are characterized by a terminal pore to the outside or the respective lumen (Slifer 1970; Falk et al. 1976; Altner and Prillinger 1980). This pore is usually formed by the cuticle tube, a structure thicker and less electron-dense than the dendritic sheath, first observed in *Musca domestica* larval sensilla (Chu-Wang and Axtell 1972) (e.g., Figure 2B, Figure 2S1-S5 E; Figure 3S1-S7 D; Figure 4S1-S2 C; Figure 4S3 D). The origin of this structure remains unclear, but most likely, it originates from the dendritic sheath and is, therefore, built by the thecogen cell.

**Figure 2.**
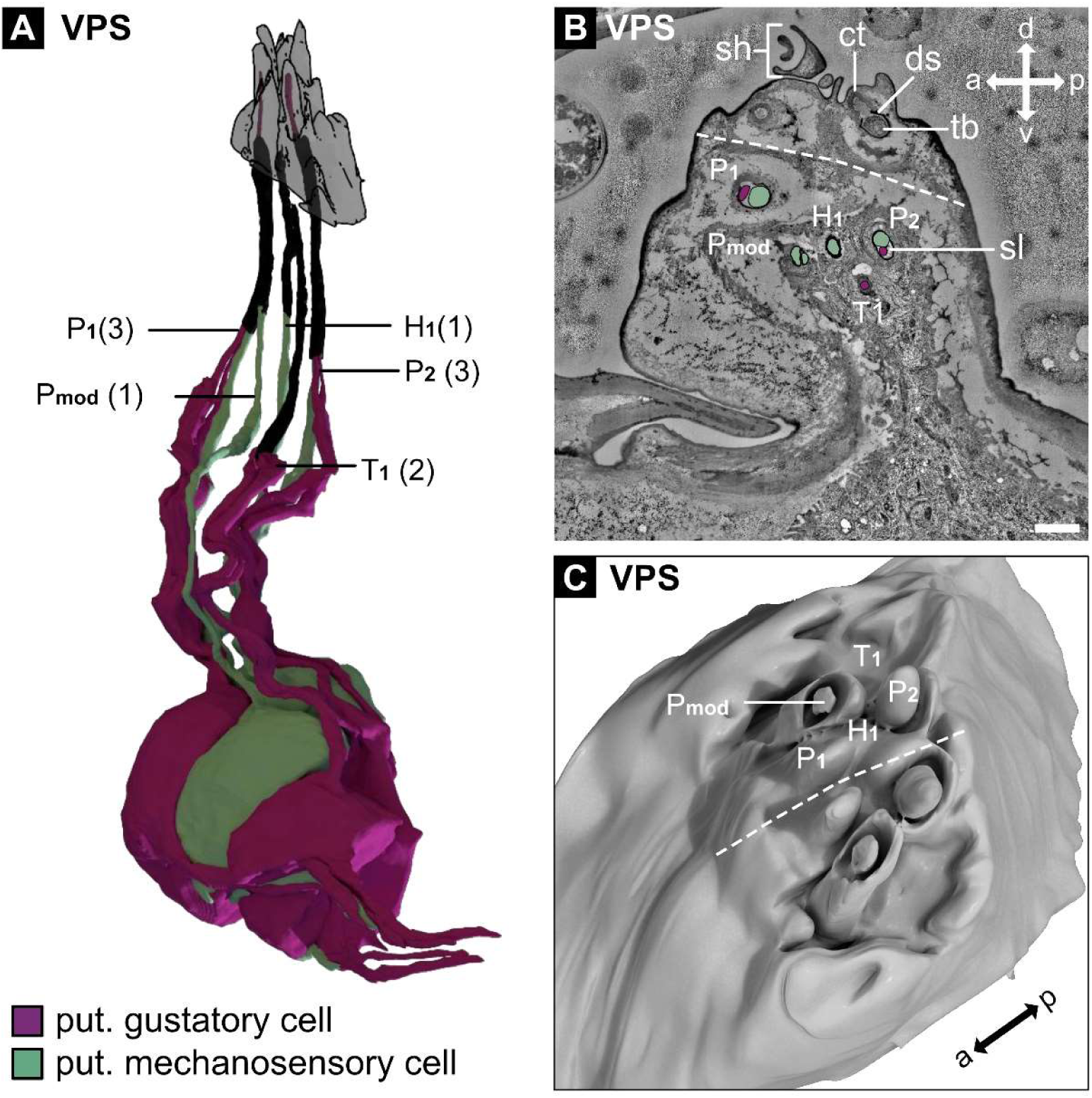
Main cellular configuration of the ventral pharyngeal sensilla (VPS). (A) 3D reconstruction of the sensory neurons innervating the VPS. The VPS is a bilaterally organized organ, but the left and right sides are fused. Only the reconstructions of the five sensilla on the left are shown. These are two papillum sensilla (P_1_ and P_2_), a modified papillum sensillum (P_mod_), a hair-like sensillum (H_1_) and a pit sensillum (T_1_). Ultrastructural features of the outer and inner morphology were used to classify the different sensilla. These sensilla resemble canonical sensilla types that can also be found in the terminal organ (see Figure 1A), the main external gustatory organ. They display different features like a terminal pore (pit sensillum - T) or a sensillum shaft (modified papillum sensillum - Pmod; hair-like sensillum - H) or both (papillum sensillum - P). Mechanosensory cells are identified by their inner dendritic morphology, which contains a tubular body, a structure known to be important for mechanosensation. For a detailed description of the single VPS sensilla, see supplementary Figures 2 S1 – S5. Nomenclature for sensilla was adapted from previous work (Chu-Wang and Axtell 1972; Rist and Thum 2017). The number of neurons innervating the sensilla is indicated in brackets. (B) STEM section of the VPS showing the organs midline (dashed line). The sensilla of the right side are highlighted, and the innervating neurons are color-coded for their putative sensory function (purple - gustatory; green - mechanosensory). (C) 3D reconstruction of the outer morphology of the VPS. The position of the sensilla is indicated for the right side of the fused organ. Unlike the other pharyngeal sensilla (see Figure 3 and Figure 4), the VPS display prominent outer structures that extend into the pharyngeal cavity. Abbreviations: d - dorsal; v - ventral; a - anterior, p - posterior, P – papillum sensillum, Pmod - papillum sensillum, H - hair sensillum, T - pit sensillum Scale bars: (B) 1 µm

**Figure 3.**
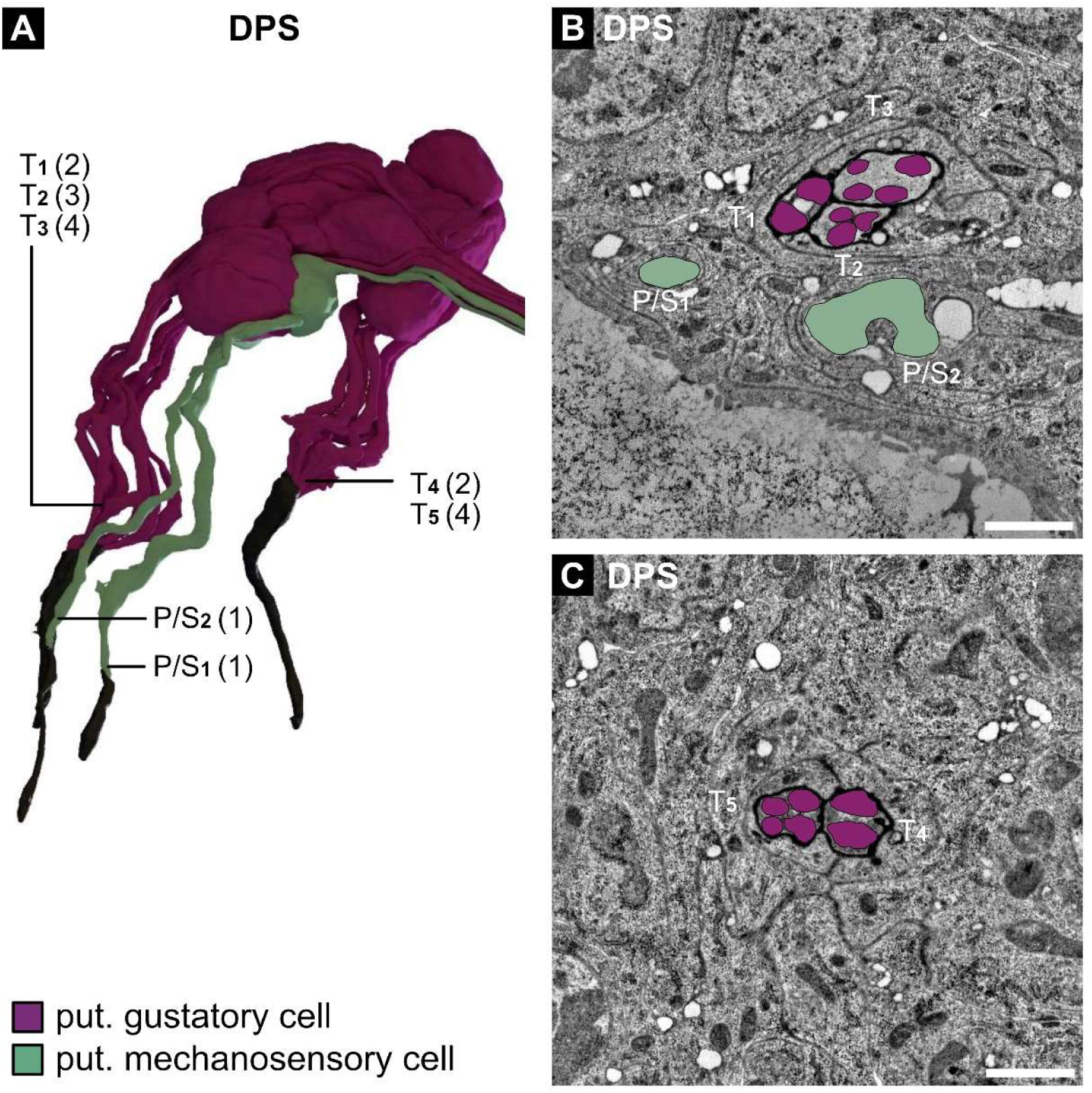
Main cellular configuration of the dorsal pharyngeal sensilla (DPS). (A) 3D reconstruction of the sensory neurons innervating the DPS. The DPS is a bilaterally organized organ; only the reconstructions of the seven sensilla on the left are shown. These are two papilla/spot sensilla (P/S_1_ and P/S_2_) and five pit sensilla (T_1_ -T_5_). From the ganglion, the sensilla expand to an anterior and a posterior location. In the anterior group, we find three pit sensilla (T_1_, T_2_ and T_3_) and two papilla/spot sensilla (P/S_1_ and P/S_2_), the latter of which is located in a deep channel. The pit sensilla display an idiosyncrasy as they share their terminal pore into the pharyngeal lumen. The posterior group consists of two pit sensilla (T_4_ and T_5_), which share a terminal pore, too. Ultrastructural features of the outer and inner morphology were used to classify the sensilla. For a detailed description of the single DPS sensilla, see supplementary Figures 3 S1 – S7. The nomenclature for sensilla was adapted from previous work (Chu-Wang and Axtell 1972; Rist and Thum 2017). The number of neurons innervating the sensilla is indicated in brackets. (B and C) STEM section of the anterior group (B) and posterior group (C) of the DPS. The sensory neurons innervating the sensilla are highlighted and color-coded for their putative sensory function (purple - gustatory; green - mechanosensory). Abbreviations: d - dorsal; v - ventral; a - anterior, p - posterior, P/S – papilla/spot sensillum, T - pit sensillum Scale bars: (B) 1 µm; (C) 1 µm

**Figure 4.**
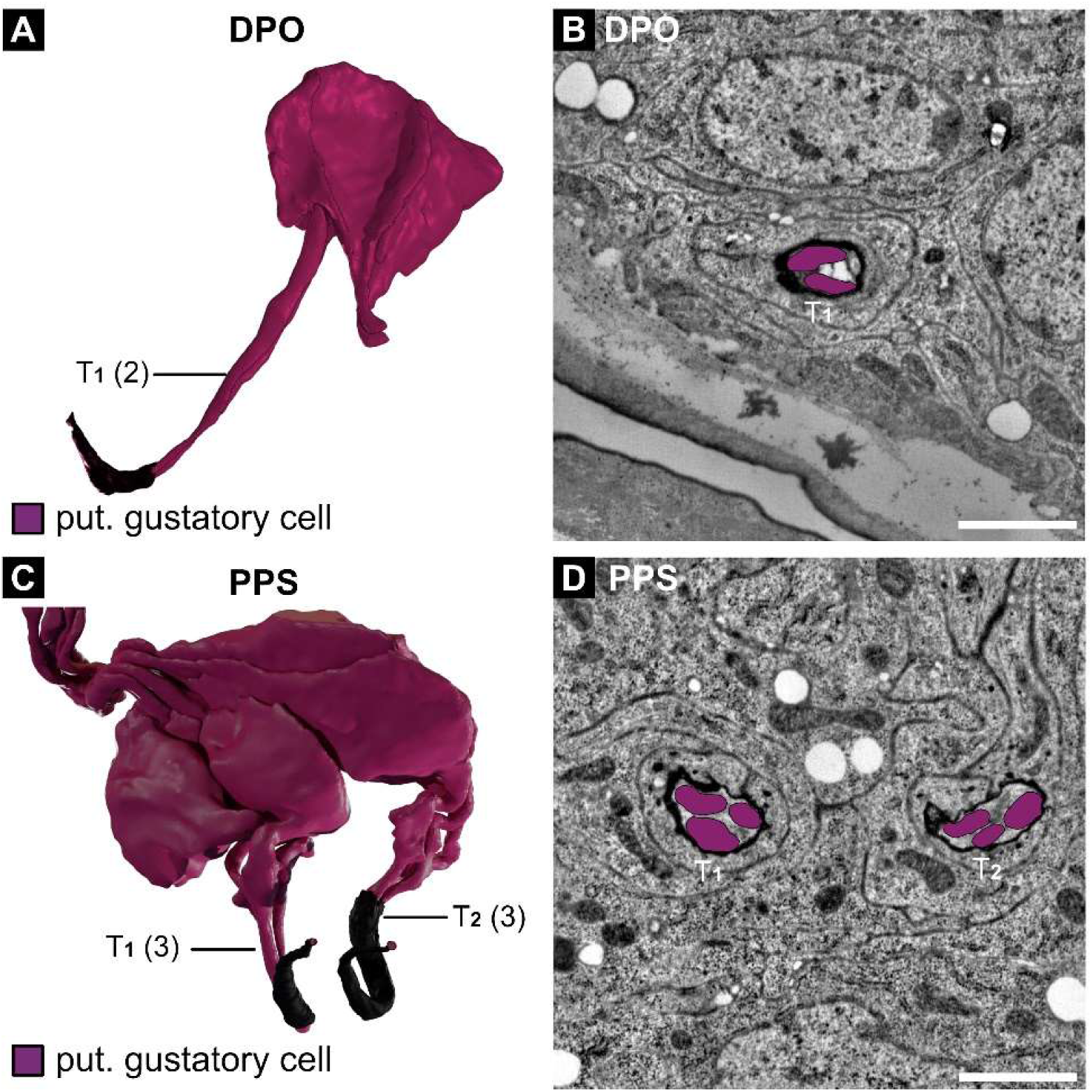
Main cellular configuration of the dorsal pharyngeal organ (DPO) and posterior pharyngeal sensilla (PPS). (A) 3D reconstruction of the sensory neurons innervating the DPO. The DPO is a bilaterally organized organ; only the reconstruction of the sensillum on the left is shown. This sensillum is a pit sensillum (T_1_) that is innervated by two sensory neurons. These are connected through a pore with the pharyngeal lumen. (B) STEM section of the DPO. The sensory neurons innervating the sensilla are highlighted and color-coded for their putative sensory function (purple - gustatory). (C) 3D reconstruction of the sensory neurons innervating the PPS. The PPS is a bilaterally organized organ; only the reconstructions of sensilla on the left are shown. These sensilla are pit sensilla (T_1_ and T_2_), innervated by three sensory neurons each. Their pores to the pharyngeal lumen are located in close proximity to each other. (D) STEM section of the PPS. The sensory neurons innervating the sensilla are highlighted and color-coded for their putative sensory function (purple - gustatory). Ultrastructural features of the outer and inner morphology were used to classify the DPO and PPS sensilla. For a detailed description of DPO and PPS sensilla, see supplementary Figures 4 S1 – S3. The nomenclature for sensilla was adapted from previous work (Chu-Wang and Axtell 1972; Rist and Thum 2017). Abbreviations: d - dorsal; v - ventral; a - anterior, p - posterior, T - pit sensillum Scale bars: (B) 1 µm; (D) 1 µm

Our analysis utilized a STEM volume of a whole first instar *Drosophila melanogaster* larva (Peale et al. 2024) to locate and reconstruct all sensory organs and their sensilla morphology (Figure 1A). While the external sensilla and the enteric sensory system were previously described using this dataset (Richter et al. 2024; Schoofs et al. 2024), we focus here on the sense organs along the pharynx, which primarily comprise gustatory sensilla that form part of the larval chemosensory system (Figure 1B). All pharyngeal organs and identified neurons are arranged bilaterally and thus organized in pairs, which we now describe in detail in the following.

### Ventral Pharyngeal Sensilla (VPS)

In contrast to other pharyngeal sensilla, the VPS displays prominent outer structures that extend into the pharyngeal lumen (Figure 2). It is a bilateral organ, but the left and right sides are fused (Figure 2A-C). Each side contains five individual sensilla innervated by a total of 10 Type I neurons per side (Figure 2A). These sensilla resemble sensilla types found in the terminal organ and other external sensilla. It consists of two papillum sensilla (P_1_ and P_2_) (Figure 2 S1-S2), one modified papillum sensillum (P_mod_) (Figure 2 S3), a hair sensillum (H_1_) (Figure 2 S4), and a pit sensillum (T_1_) (Figure 2 S5). All VPS sensilla display the standard set of support cells that enclose the sensory neurons and the sensillum lymph (Figure 2 S1-S5 F; Figure 2 S5 E), but the thecogen cells show different degrees of lamellation (Table 1; Figure 2 S1-S2 H; Figure 2 S3-S5 G). The papillum sensilla P_1_ and P_2_ display a shaft with a terminal pore to the pharyngeal lumen (Figure 2 S1-S2 C) and a cuticle tube (ct) (Figure 2 S1-S2 D) that connects with the dendritic sheath (Figure 2 S1-S2 E). Both are innervated by three dendrites each, two that protrude into the pore and, therefore, most likely serve a gustatory function. The third one ends with a tubular body (Figure 2 S1-S2 E) and, therefore, most likely, serves a mechanosensory function. P_mod_ displays a shaft that forms a cylindrical portion that encircles a bud-like structure (Figure 2 S3C). It has a single dendrite that ends with a tubular body below the bud. The dendritic sheath remains intact, and a pore is absent. P_mod_ most likely serves a mechanosensory function. H_1_ is internally built in a similar configuration as the P_mod_, containing a tubular body and no pore to the lumen. Only the sensillum shaft is built differently (Figure 2 S4C) and resembles the hair sensilla found in the larval body segments. The different outer morphologies may induce different mechanisms in mechanotransduction or, at least, different stimulus strengths needed (Figure 2 S3-S4 D, E). T_1_ resembles a pit sensillum, containing a pore to the lumen and a cuticle tube that connects to the dendritic sheath (Figure 2 S5C-D). The pore channel is innervated by two, most likely gustatory neurons. All cell bodies of neurons innervating the VPS lie in the VPS ganglion (VPSG) (Figure 2 S1-S5I). Furthermore, three multidendritic Type II neurons are present in the VPSG (Figure 2 S3I). All VPSG neurons send axons to the subesophageal zone (SEZ) via the labial nerve (LbN).

### Dorsal Pharyngeal Sensilla (DPS)

The DPS is a bilaterally organized organ that contains seven sensilla on each side, innervated by a total of 17 Type I neurons per side. From the ganglion, the neurons extend their dendrites toward anterior and posterior locations in the pharyngeal cavity (Figure 3 A-C; Figure 3S1-S7 A, B). In the anterior location, we found three pit sensilla (T_1_, T_2_, and T_3_) and two papilla/spot sensilla (P/S_1_ and P/S_2_), the latter located in a deep depression (Figure 3 S2 B). Papilla and spot sensilla display the same morphology, but the former are termed spot sensilla when found within a compound sensillum (Rist and Thum 2017) and papilla sensilla when occurring as a single sensillum (Dambly-Chaudière and Ghysen 1986; Richter et al. 2024). In the posterior location, we identified two pit sensilla (T_4_ and T_5_). In contrast to the pit sensilla of other larval gustatory organs, the DPS pit sensilla of the anterior group (T_1_, T_2_, and T_3_) and the posterior group (T_4_ and T_5_) share a cuticle tube and a terminal pore that opens into the pharyngeal lumen (Figure 3 S3-S7 B-C).

Nevertheless, each DPS sensillum possesses its own set of support cells and its own dendritic sheath that separates the dendrites of the sensilla from each other and creates an individual sensillum lymph cavity (Figure 3 S3-S7 D-F), but for T1-T3 and T4 + T5, all trichogen cells generate the respective common cuticle tube. This finding is in contrast to the proposed organization of the LSO (Nayak and Singh 1983), which suggests that the eight sensory cells are accompanied by a shared set of three support cells. These differences may arise from the superior detail and completeness of a whole volume EM stack compared to conventional EM. The thecogen cells display varying degrees of lamellation, ranging from absent in P/S_1_ and P/S_2_, to weak in T_1_, T_2_, and T_3_, and pronounced in T_4_ and T_5_ (Figure 3 S1-S7 E-F). All pit sensilla neurons are most likely gustatory, as their dendrite endings are in contact with the pharyngeal lumen and do not display another morphological feature indicating a different sensory modality. The DPS papilla/spot sensilla most likely serve a mechanosensory function, as their dendrite tips contain a tubular body, and the dendritic sheath does not merge into a cuticle tube (Figure 3 S1-S2 C-D). Most somata of neurons innervating the DPS lie in the DPS ganglion (DPSG) (Figure 3 S2-S7 G). Only the soma of P/S_1_ lies nearby outside the ganglion (Figure 3 S1 G). All DPS neurons send their axons to the subesophageal zone (SEZ) via the labral nerve (LrN).

### Dorsal Pharyngeal Organ (DPO)

The DPO is a small, bilateral, organized structure that contains only one pit sensillum (T_1_) on each side, innervated by two Type I neurons (Figure 4 A-B; Figure 4S1 A). From the ganglion, the dendrites extend toward a location posterior to the DPS. The pit sensillum features a cuticle tube (Figure 4S1 B) and a terminal pore (Figure 4S1 C) leading to the pharyngeal lumen. Therefore, the two innervating neurons are most likely gustatory. The sensillum includes the standard repertoire of support cells that enclose the sensory neurons and the sensillum lymph (Figure 4S1 D-E). The thecogen cell exhibits weak lamellations (Figure 4S1 F). All cell bodies of neurons innervating the DPO are located in the DPO ganglion (DPOG).

Additionally, three multidendritic Type II neurons, here named MD1, MD2, and MD3, are located in the DPOG (Figure 4 S1G to I). These neurons extend multiple dendrites into the extracellular space, where, based on their position and morphology, they are likely to perceive internal chemosensory, thermosensory, or mechanosensory signals. The dendrites of MD1 and MD2 wrap around the cell body of MD3 and form a bulbous structure in which multiple dendrites branch and intertwine. This structure resembles mechanoreceptors in human skin, such as Pacinian, Meissner, or Krause corpuscles, and may serve a similar mechanosensory function, although chemosensory or thermosensory roles are also possible (Krause 1880; Paré et al. 2001; Handler and Ginty 2021; Ziolkowski et al. 2025). In contrast, the dendrites of MD3 extend toward nearby muscle cells, suggesting a potential role in sensing muscle movements during ingestion, while a chemosensory function cannot be excluded. The markedly different dendritic arborizations of MD1, MD2, and MD3 suggest that these neurons represent distinct sensory types that may be tuned to different modalities, sensitivities, or stimulus directionalities. Although their precise functional roles remain unclear, we propose that these neurons act as internal sensors detecting hemolymph signals and or mechanical cues associated with muscle activity. All DPOG neurons project their axons to the subesophageal zone via the labral nerve.

### Posterior Pharyngeal Sensilla (PPS)

The PPS is a bilateral organized structure containing two pit sensilla (T_1_ and T_2_) on each side that are innervated by three Type I neurons each (Figure 4C-D). From the ganglion, the dendrites extend towards the posterior-most location in the pharynx, right before the esophagus (Figure 1B). The pit sensilla display a cuticle tube and a terminal pore to the pharyngeal lumen. Therefore, the three innervating neurons are most likely gustatory. All cell bodies of neurons innervating the PPS lie in the PPS ganglion (PPSG) (Figure 4 S2-S3). All PPSG neurons send their axons to the subesophageal zone (SEZ) via the labral nerve (LrN).

### Additional sensory structures

In addition to the main pharyngeal sense organs, we find three single papilla/spot sensilla associated with the feeding apparatus. These resemble classic mechanosensory sensilla found in the larval segments. They possess a tubular body at their dendritic tip, and a dendritic sheath encloses the dendrite. Their dendrites protrude towards the pharyngeal lumen anterior to the DPS/VPS region, and their axons run with the labial nerve (LbN) (Figure 4S4 A-D).

Furthermore, four chordotonal organs are also associated with the larval feeding apparatus (Richter et al. 2024). Three lie along the ventral part of the cephalopharyngeal skeleton (CPS), with the anterior two being monodynal (innervated by a single sensory neuron) and the posterior one being heterodynal (innervated by two sensory neurons). Their axons run with the labial nerve (LbN). The fourth one can be found in the dorsal posterior portion of the CPS. It is also monodynal, and its axon runs along the labral nerve (LN) (Figure 4S4 A, E-H).

In addition to the Type I sensory neurons organized in the sensilla, we also identified six multidendritic neurons associated with the larval feeding apparatus. As mentioned above, three are located within the VPS ganglion, and three are found in the DPO ganglion.

### Summary/Conclusion

The larval pharyngeal sensory system comprises four major organs containing 15 sensilla (four mechanosensory, nine chemosensory, two dual-function), with 35 Type I neurons (six mechanosensory, 29 chemosensory) and six Type II neurons in total. Additional sensory structures include three papilla/spot sensilla and four chordotonal organs, distributed across the cephalopharyngeal region, completing our understanding of the larval sensory system. For the first time, the exact sensilla and cell numbers for the pharyngeal organs and sensilla have been described, and a functional prediction of their sensory modality has been made.

In total, 124 Type I neurons are running with the major head nerves (MxN – 47; LbN – 14; AN – 37; LrN – 26). Based on our and previous analyses, there are 55 gustatory or chemosensory neurons, 40 mechanosensory or proprioceptive neurons, 21 olfactory neurons, six thermosensory neurons, and two neurons with unidentified function (for details, see Table 2). We conclude that these neurons, in conjunction with the enteric system, constitute the primary sensory apparatus for making feeding decisions. This includes navigating towards desirable conditions, evaluating the chemical and physical quality of food, and incorporating this information with the internal state of the animal.

**Table 2.**
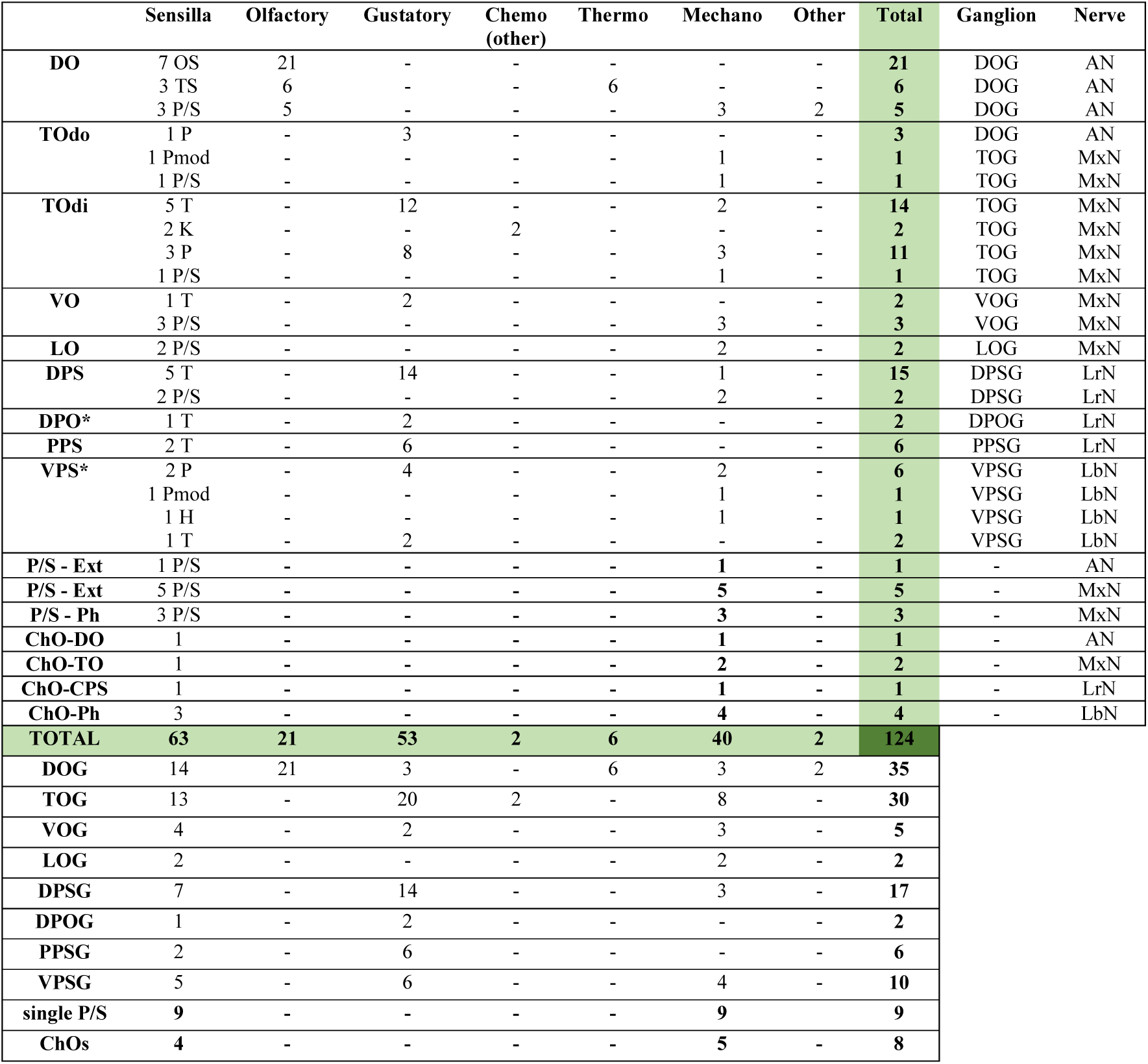
Neuronal composition and putative modality of larval head organs and their nerves. Table updated from Python and Stocker (2002). This summary presents a synopsis of previously reported cell numbers and identities (Singh and Singh 1984; Campos-Ortega and Hartenstein 1985; Schmidt-Ott et al. 1994; Rist and Thum 2017; Richter et al. 2024), as well as the data presented in this work. * In the VPS and DPS ganglia, there are three cell bodies of multidendritic neurons each, which in sum is in accordance with previously reported cell numbers. Multidendritic neurons (MDNs) have generally not been considered here; however, based on previously reported cell numbers in the head nerves (Miroschnikow et al., 2018), we propose that there are approximately 25 sensory MDNs (per side) innervating the head region through the MxN, AN, and PaN. Additionally, there are 36 sensory MDNs (in total) projecting through the larval vagus nerve (VN) and innervating the aorta, esophagus and gut. Abbreviations: DO – dorsal organ; TO – terminal organ; TOdo – dorsal group of the terminal organ; TOdi – distal group of the terminal organ; VO – ventral organ; LO – labial organ; DPS – dorsal pharyngeal sensilla; DPO – dorsal pharyngeal organ; PPS – posterior pharyngeal sensilla; VPS – ventral pharyngeal sensilla; P/S – papilla/spot sensillum; ChO – chordotonal organ; OS – olfactory sensillum; TS – thermosensory sensillum; P – papillum sensillum; Pmod – modified papillum sensillum; T – pit sensillum; K – knob sensillum; H – hair sensillum; DO - dorsal organ ganglion; TOG - terminal organ ganglion; VOG - ventral organ ganglion; LOG - labial organ ganglion; DPSG – dorsal pharyngeal sensilla ganglion; DPOG – dorsal pharyngeal organ ganglion; PPSG – posterior pharyngeal sensilla ganglion; VPSG – ventral pharyngeal sensilla ganglion; CPS – cephalopharyngeal skeleton; Ph – Pharynx; Ext – external; AN – antennal nerve; MxN – maxillary nerve; LrN – labral nerve; LbN – labial nerve

|  | Sensilla | Olfactory | Gustatory | Chemo<br>(other) | Thermo | Mechano | Other | Total | Ganglion | Nerve |
| --- | --- | --- | --- | --- | --- | --- | --- | --- | --- | --- |
| **DO** | 7 OS | 21 | - | - | - | - | - | **21** | DOG | AN |
|  | 3 TS | 6 | - | - | 6 | - | - | **6** | DOG | AN |
|  | 3 P/S | 5 | - | - | - | 3 | 2 | **5** | DOG | AN |
| **TOdo** | 1 P | - | 3 | - | - | - | - | **3** | DOG | AN |
|  | 1 Pmod | - | - | - | - | 1 | - | **1** | TOG | MxN |
|  | 1 P/S | - | - | - | - | 1 | - | **1** | TOG | MxN |
| **TOdi** | 5 T | - | 12 | - | - | 2 | - | **14** | TOG | MxN |
|  | 2 K | - | - | 2 | - | - | - | **2** | TOG | MxN |
|  | 3 P | - | 8 | - | - | 3 | - | **11** | TOG | MxN |
|  | 1 P/S | - | - | - | - | 1 | - | **1** | TOG | MxN |
| **VO** | 1 T | - | 2 | - | - | - | - | **2** | VOG | MxN |
|  | 3 P/S | - | - | - | - | 3 | - | **3** | VOG | MxN |
| **LO** | 2 P/S | - | - | - | - | 2 | - | **2** | LOG | MxN |
| **DPS** | 5 T | - | 14 | - | - | 1 | - | **15** | DPSG | LrN |
|  | 2 P/S | - | - | - | - | 2 | - | **2** | DPSG | LrN |
| **DPO\*** | 1 T | - | 2 | - | - | - | - | **2** | DPOG | LrN |
| **PPS** | 2 T | - | 6 | - | - | - | - | **6** | PPSG | LrN |
| **VPS\*** | 2 P | - | 4 | - | - | 2 | - | **6** | VPSG | LbN |
|  | 1 Pmod | - | - | - | - | 1 | - | **1** | VPSG | LbN |
|  | 1 H | - | - | - | - | 1 | - | **1** | VPSG | LbN |
|  | 1 T | - | 2 | - | - | - | - | **2** | VPSG | LbN |
| **P/S - Ext** | 1 P/S | - | - | - | - | 1 | - | **1** | - | AN |
| **P/S - Ext** | 5 P/S | - | - | - | - | 5 | - | **5** | - | MxN |
| **P/S - Ph** | 3 P/S | - | - | - | - | 3 | - | **3** | - | MxN |
| **ChO-DO** | 1 | - | - | - | - | 1 | - | **1** | - | AN |
| **ChO-TO** | 1 | - | - | - | - | 2 | - | **2** | - | MxN |
| **ChO-CPS** | 1 | - | - | - | - | 1 | - | **1** | - | LrN |
| **ChO-Ph** | 3 | - | - | - | - | 4 | - | **4** | - | LbN |
| **TOTAL** | **63** | **21** | **53** | **2** | **6** | **40** | **2** | **124** |  |  |
| **DOG** | 14 | 21 | 3 | - | 6 | 3 | 2 | **35** |  |  |
| **TOG** | 13 | - | 20 | 2 | - | 8 | - | **30** |  |  |
| **VOG** | 4 | - | 2 | - | - | 3 | - | **5** |  |  |
| **LOG** | 2 | - | - | - | - | 2 | - | **2** |  |  |
| **DPSG** | 7 | - | 14 | - | - | 3 | - | **17** |  |  |
| **DPOG** | 1 | - | 2 | - | - | - | - | **2** |  |  |
| **PPSG** | 2 | - | 6 | - | - | - | - | **6** |  |  |
| **VPSG** | 5 | - | 6 | - | - | 4 | - | **10** |  |  |
| **single P/S** | 9 | - | - | - | - | 9 | - | **9** |  |  |
| **ChOs** | 4 | - | - | - | - | 5 | - | **8** |  |  |

Future work will be needed to trace the peripheral sensory system of the larval pharynx from the sensory organs to the brain via the axons of its associated sensory neurons, and to align their central terminals with those identified in previous studies, including Miroschnikow et al. (2018). Such analyses will provide a basis for defining the sensory subcircuits of the larval feeding network and for understanding how their wiring principles transform food-related information into appropriate behavioral outputs.

In conclusion, this comprehensive characterization of the *Drosophila* larval pharyngeal sensory system provides an unprecedented mapping of neuronal architecture and function. This detailed neuroanatomical foundation will help to gain new insights into sensorimotor integration, decision-making circuits, and the neural basis of adaptive behaviors in response to environmental changes.

## Materials and Methods

### Whole larval volume

Sensilla and neuron reconstruction were performed on a STEM (scanning transmission electron microscopy) volume of a whole first-instar larva; technical details of its generation are provided in Schoofs et al. (2024) and Peale et al. (2024). Briefly, a first instar larva was sectioned into 4,866 sections of 34 nm thickness that were placed on 1622 grids (3 per grid), using a system for automated section cutting and pickup called iTome (short for interferometric microtome). For imaging of the sections, the grids were post-stained with uranyl acetate and lead acetate and then placed into a Zeiss Ultra 55 (modified for operation as a STEM). The micrographs were then reassembled into one dataset (Peale et al. 2024).

We identified sensory structures by scanning the dataset in the pharyngeal region, thereby recognizing dendritic processes towards the pharyngeal lumen. Localization of all sensilla and neurons in the pharyngeal region was done using CATMAID (http://www.catmaid.org (Saalfeld et al. 2009)).

### Image processing

We extracted smaller volumes of the sensory organs from the entire larval volume. The obtained image stacks were imported into Amira (Thermo Fischer Scientific, v2019). Since the whole larval volume was already aligned, the stacks were only slightly realigned by manual correction. In Amira, structures of interest were segmented and transformed into 3D objects. Next, the segmentations were imported to Blender (Blender Institute, Amsterdam), where the 3D reconstructions were manually finished using the preliminary segmentation as a template.

## Acknowledgements

This work was supported by the Deutsche Forschungsgemeinschaft (Grant No. 441181781, 426722269, 432195391), EU funds from the ESF Plus Program (Grant No. 100649752), and the Open Access Publishing Fund of Leipzig University, which is supported by the German Research Foundation within the Open Access Publication Funding program.

We thank Michael Pankratz, Anton Miroschnikow, Andreas Schoofs, Wolf Hütteroth, Katharina Eichler, and the members of the behavioral neurogenetics group for their support, help, discussions, and comments. We also thank Philip Schlegel, Casey Schneider-Mizell, and Tom Kazimiers for assistance with the whole larva STEM volume.

## Data availability statement

The data supporting this study’s findings are available from the corresponding author upon reasonable request. The full larval EM volume was published in Peale et al, 2024.

