## Supplemental figures and tables for "Morphology and ultrastructure of pharyngeal sense organs of *Drosophila* larvae"

**Supplementary table 1: Comparison of pharyngeal and cibarial organs in different insect species.**

| Taxon | Species | organ name | sensillum type | sensilla | neurons | sensory modality | references |
|---|---|---|---|---|---|---|---|
| ***DIPTERA*** | *D. melanogaster* (larvae) | DPS | papilla/spot sensilla, pit sensilla | 7 | 17 | Mechanosensory; Gustatory | Gendre 2004; this work |
| | | DPO | pit sensilla | 1 | 2 (+3*) | Gustatory | Gendre 2004; this work |
| | | VPS | pit sensilla, papillum sensilla, hair sensilla | 5 | 10 | Mechanosensory; Gustatory | Gendre 2004; this work |
| | | PPS | pit sensilla | 2 | 6 | Gustatory | Gendre 2004; this work |
| | | single sensilla | papilla/spot sensilla | 3 | 3 | Mechanosensory | Gendre 2004; this work |
| | *D. melanogaster* (adult) | LSO | bristle sensilla, hair sensilla, hairless sensilla | 9 | 18 | Mechanosensory; Gustatory | Nayak and Singh 1983; Gendre 2004 |
| | | VCSO | bristle sensilla, hair sensilla, hairless sensilla | 3 | 8 | Gustatory | Nayak and Singh 1983; Gendre 2004 |
| | | DCSO | bristle sensilla, hair sensilla, hairless sensilla | 2 | 6 | Gustatory | Nayak and Singh 1983; Gendre 2004 |
| | | ventral and dorsal row sensilla | fishtrap bristle sensilla | 18-23 | 1 | Mechanosensory | Nayak and Singh 1983; Gendre 2004 |
| | *C. vicinia (*larvae) | LbO (=LSO) | N/A | N/A | N/A | Mechanosensory | Ludwig 1949; Hückesfeld 2010 |
| | *C. vicinia* (adult) | CO | basiconic, trichoid and campaniform sensilla | 24 | N/A | Mechanosensory; Gustatory | Rice 1973 |
| | *M. domestica* (larva) | LSO | N/A | N/A | N/A | Mechanosensory; Gustatory | Ranade, 1967 |
| | *A. aegypti (*adult) | Cibarial Sensilla | Papilla sensilla, campaniform sensilla | 9 (female), 7 (male) | N/A | Mechanosensory; Gustatory | McIver 1981; Lee 1983; Baik 2020 |
| ***LEPIDOPTERA*** | *M. sexta* (larvae) | Epipharyngeal sensilla | N/A | 1 | 3 | Gustatory | Glendinning 2000 |
| | *Heterogynis penella* (larvae) | Cibarial sensilla | N/A | N/A | N/A | N/A | Vegliante 2005 |
| ***COLEOPTERA*** | *L. decemlineata* (larvae) | Epipharyngeal sensilla | N/A | 10 | N/A | Gustatory | Messchendorp 1998 |
| ***HEMIPTERA*** | *R. proxilus* (adult) | Pharyngeal Organ (PO) | Short-peg sensilla | 8** | N/A | Gustatory | Ortega-Insaurralde 2024 |
| ***HYMENOPTERA*** | *A. mellifera* (adult) | Epipharyngeal sensilla | N/A | ~180 | N/A | Gustatory | Galic 1971 |
| ***HYMENOPTERA*** | *A. mellifera* (adult) | Hypopharyngeal sensilla | N/A | ~90 | N/A | Gustatory | Galic 1971 |
| ***ORTHOPTERA*** | *Acheta domesticus* (adult) | Pharyngeal neurons | N/A | N/A | N/A | N/A | Kirby 1984 |
| ***ODONATA*** | *Ischnura elegans (adult)* | Epipharyngeal sensilla | hair sensilla, peg sensilla | 30** (hair); 50** (pegs) | N/A | Mechanosensory; Gustatory | Rebora 2014 |

****multidendritic neurons; ** unpaired***

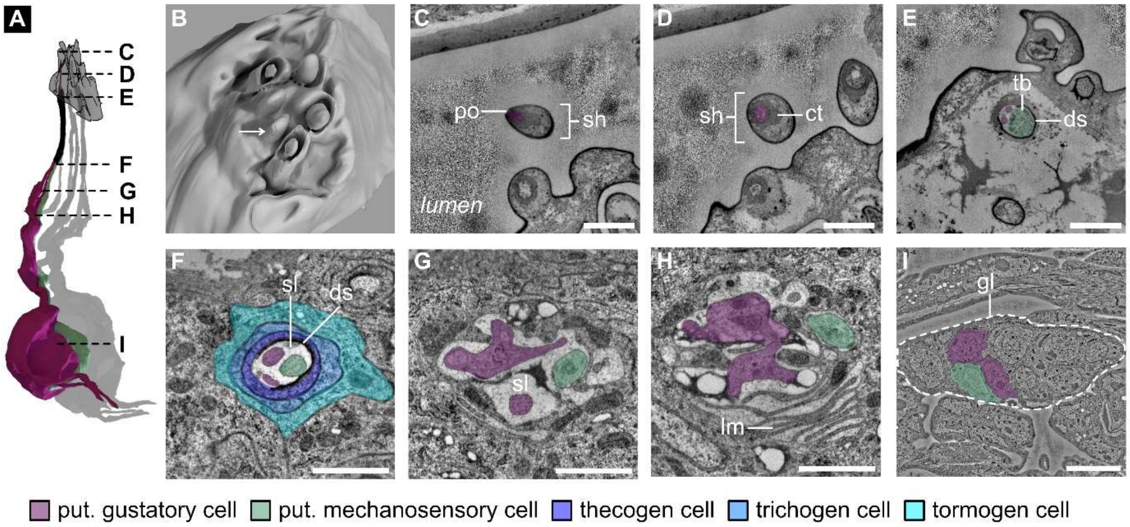


**Figure 2S1 – VPS-P$_1$. Internal morphology of the papillum sensillum 1 (P$_1$) of the ventral pharyngeal sensilla (VPS).**

(A) 3D reconstruction of the sensory neurons innervating the P$_1$ sensillum. Three sensory neurons innervate P$_1$. (B) 3D reconstruction of the outer morphology of the VPS. The arrow indicates the location of the P$_1$ sensillum. (C-I) Serial STEM sections of the P$_1$ sensillum. (C and D) The sensillum shaft is visible. The terminal pore is innervated by sensory neurons (purple). (E) Further proximal, all three cilia are visible. Two neurons (purple) are innervating the terminal pore, and one neuron (turquoise) ends at the base of the pore with a tubular body. A dendritic sheath is present. (F) The three dendrites are bathed in the sensillum lymph and enclosed by the dendritic sheath. The thecogen, trichogen and tormogen support cells are highlighted. (G) The three dendrites are bathed in the sensillum lymph, but the dendritic sheath is absent at this level. (H) The sheath cell is highly compartmented and lamellated at this level. (I) Section of the VPS ganglion. The cell bodies of the three P$_1$ neurons are visible.

Abbreviations: po - pore; sh - shaft; ct - cuticle tube; ds - dendritic sheath; tb - tubular body; sl - sensillum lymph; lm - lamellae

Scale bars: C-H: 1 µm; I: 5 µm

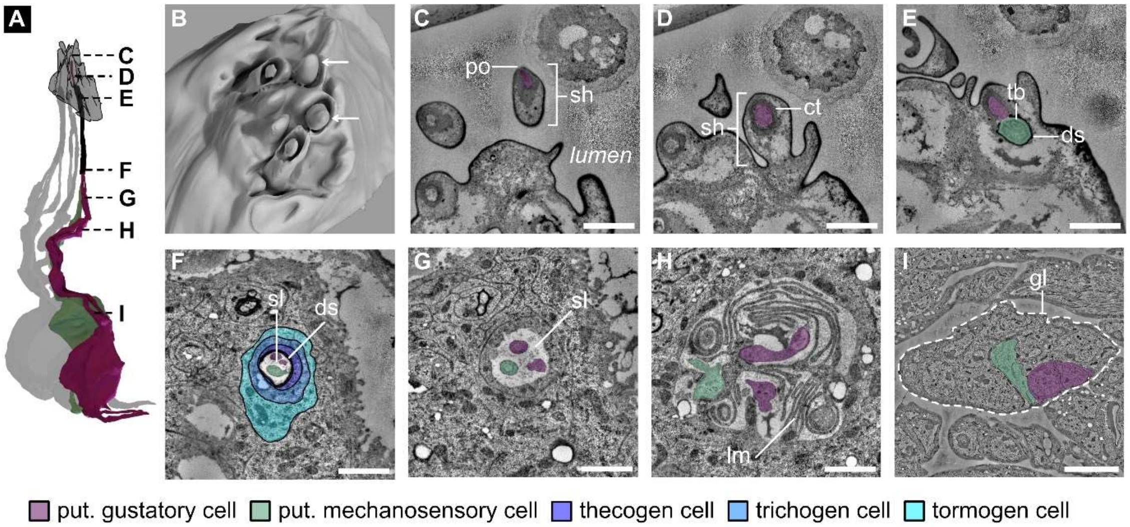


**Figure 2S2 – VPS-$P_2$. Internal morphology of the papillum sensillum 2 ($P_2$) of the ventral pharyngeal sensilla (VPS).**

(A) 3D reconstruction of the sensory neurons innervating the $P_2$ sensillum. Three sensory neurons innervate $P_2$. (B) 3D reconstruction of the outer morphology of the VPS. The arrow indicates the location of the $P_2$ sensillum. (C-I) Serial STEM sections of the $P_2$ sensillum. (C and D) The sensillum shaft is visible. The terminal pore is innervated by sensory neurons (purple). (E) Further proximal, all three cilia are visible. Two neurons (purple) are innervating the terminal pore, and one neuron (turquoise) ends at the base of the pore with a tubular body. A dendritic sheath is present. (F) The three dendrites are bathed in the sensillum lymph and enclosed by the dendritic sheath. The thecogen, trichogen and tormogen support cells are highlighted. (G) The three dendrites are bathed in the sensillum lymph, but the dendritic sheath is absent at this level. (H) The sheath cell is highly compartmented and lamellated at this level. (I) Section of the VPS ganglion. The cell bodies of the three $P_2$ neurons are visible.

Abbreviations: po - pore; sh - shaft; ct - cuticle tube; ds - dendritic sheath; tb - tubular body; sl - sensillum lymph; lm - lamellae; gl - ganglion

Scale bars: C-H: 1 µm; I: 5 µm

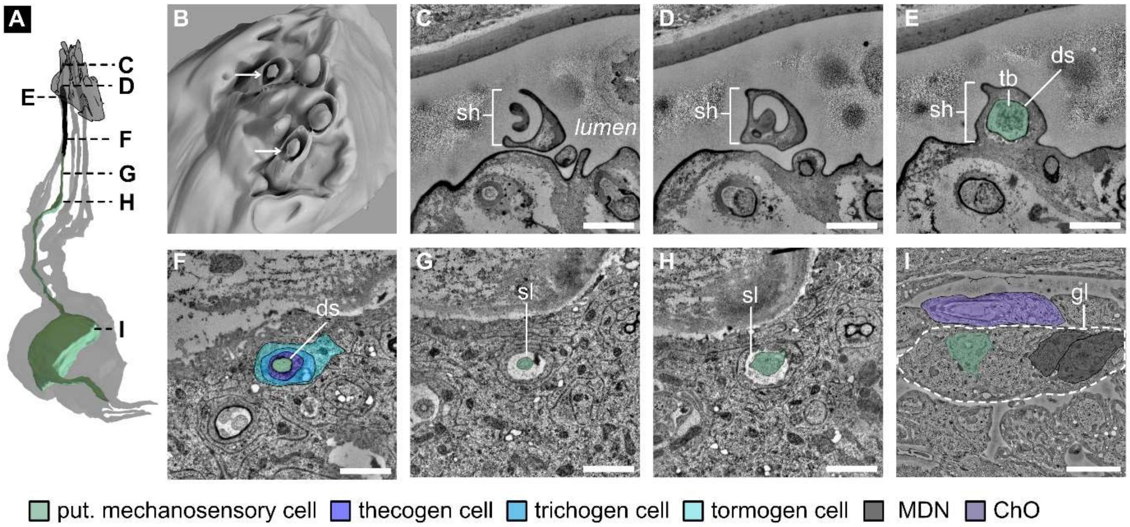


**Figure 2S3 – VPS-$P_{mod}$. Internal morphology of the modified papillum sensillum ($P_{mod}$) of the ventral pharyngeal sensilla (VPS).**

(A) 3D reconstruction of the $P_{mod}$ sensillum. A single sensory neuron innervates $P_{mod}$. (B) 3D reconstruction of the outer morphology of the VPS. The arrow indicates the location of the $P_{mod}$ sensillum. (C-I) Serial STEM sections of the $P_{mod}$ sensillum. (C and D) The sensillum shaft is visible. Its shaft forms a cylindrical portion that encircles a bud-like structure. The bud lacks a terminal pore. (E) Further proximal, the neuron (turquoise) at the base of the shaft is visible. The neuron ends with a tubular body. A dendritic sheath is present. (F) The dendrite is entirely enclosed by the dendritic sheath. The thecogen, trichogen and tormogen support cells are highlighted. (G and H) The dendrite is bathed in the sensillum lymph, but the dendritic sheath is absent at this level. The sheath cell is not compartmented and lamellated. (I) Section of the VPS ganglion (VPSG). The cell body of the $P_{mod}$ neuron is visible. Additionally, three cell bodies of multidendritic sensory neurons (Type II neurons) lie in the VPSG (two cell bodies and one dendrite visible in this section). A heterodynal chordotonal organ lies close to the VPSG.

Abbreviations: sh – shaft; tb – tubular body; ds – dendritic sheath; sl – sensillum lymph; gl – ganglion; MDN – multidendritic neuron; ChO – chordotonal organ

Scale bars: C-H: 1 µm; I: 5 µm

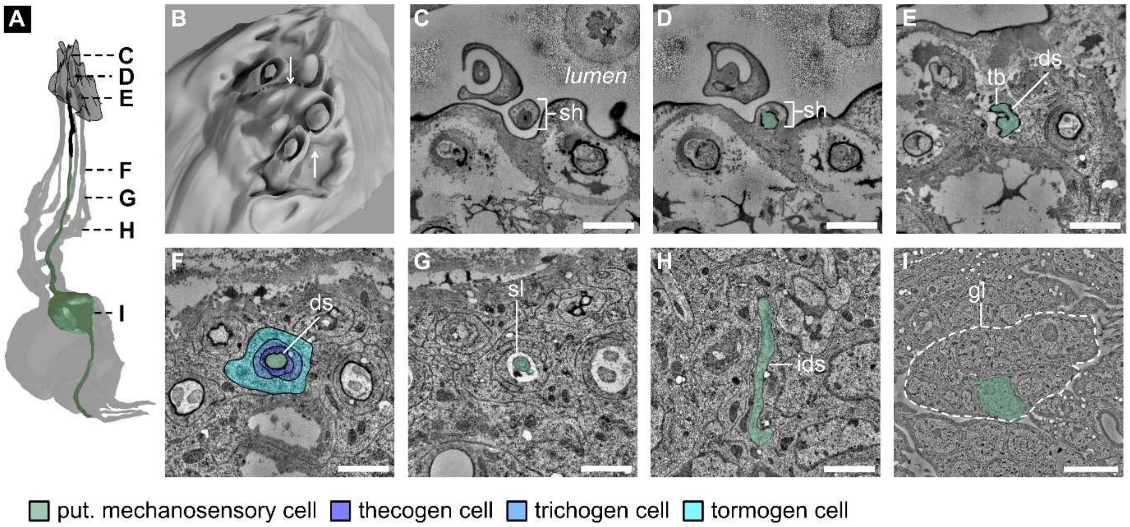


**Figure 2S4 – VPS-$H_1$. Internal morphology of the hair-like sensillum 1 ($H_1$) of the ventral pharyngeal sensilla (VPS).**

(A) 3D reconstruction of the $H_1$ sensillum. A single sensory neuron innervates $H_1$. (B) 3D reconstruction of the outer morphology of the VPS. The arrow indicates the location of the $H_1$ sensillum. (C-I) Serial STEM sections of the $H_1$ sensillum. (C and D) The sensillum shaft is visible. Its shaft forms a hair-like structure. The hair lacks a terminal pore. Externally, the shaft is hardly visible as the other prominent VPS sensilla mask it. (D and E) The neuron (turquoise) at the base of the shaft is visible. The neuron ends with a tubular body. A dendritic sheath is present. (F) The dendrite is entirely enclosed by the dendritic sheath. (G) The dendrite is bathed in the sensillum lymph, but the dendritic sheath is absent at this level. The thecogen, trichogen and tormogen support cells are highlighted. (H) The sheath cell is not compartmented and lamellated. (I) Section of the VPS ganglion. The cell body of the $H_1$ neuron is visible.

Abbreviations: sh - shaft; ds - dendritic sheath; tb - tubular body; sl - sensillum lymph; ids - inner dendritic segment; gl - ganglion

Scale bars: C-H: 1 µm; I: 5 µm

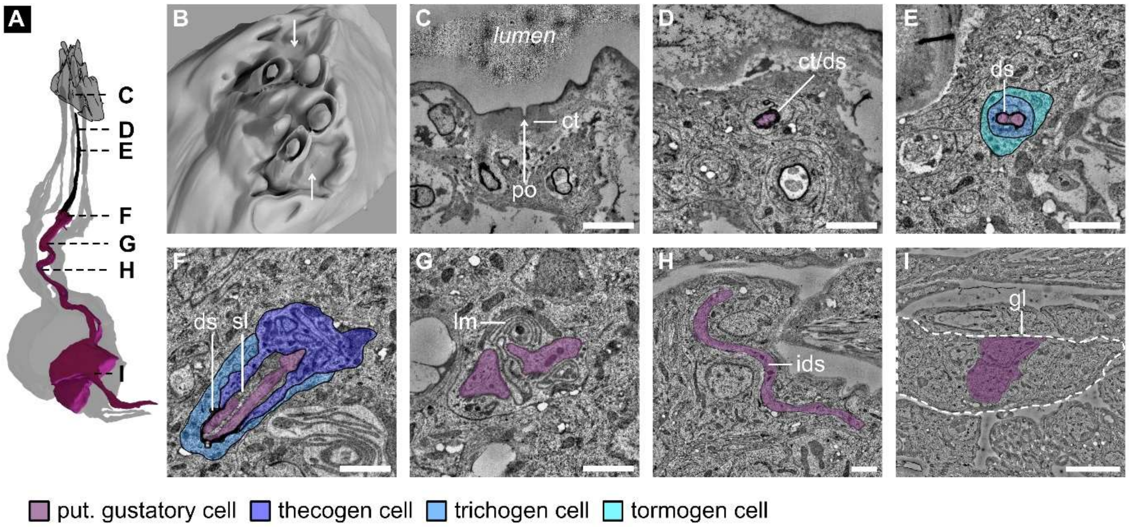


**Figure 2S5 – VPS-$T_1$. Internal morphology of the pit sensillum 1 ($T_1$) of the ventral pharyngeal sensilla (VPS).**

(A) 3D reconstruction of the $T_1$ sensillum. Two sensory neurons innervate $T_1$. (B) 3D reconstruction of the outer morphology of the VPS. The arrow indicates the location of the $T_1$ sensillum. (C-I) Serial STEM sections of the $T_1$ sensillum. (C) The terminal pore is visible. A cuticle tube connects to the lumen through a pore and is innervated by the dendrites. (D) The cuticle tube connects with the dendritic sheath. (E) The two ciliary structures (purple) are visible. A dendritic sheath is present. The trichogen and tormogen support cells are highlighted. (F) The tips of the dendrites are enclosed by the dendritic sheath, and the proximal parts are bathed in the sensillum lymph enclosed by the sheath cell. The thecogen and trichogen support cells are highlighted. (G) The dendrite is bathed in the sensillum lymph, but the dendritic sheath is absent at this level. The sheath cell is lamellated. (H) Inner dendritic segment of one sensory cell (I) Section of the VPS ganglion. The cell bodies of the $T_1$ neurons are visible.

Abbreviations: po - pore; ct - cuticle tube; ds - dendritic sheath; sl - sensillum lymph; lm - lamellae; ids - inner dendritic segment; gl - ganglion

Scale bars: C-H: 1 µm; I: 5 µm

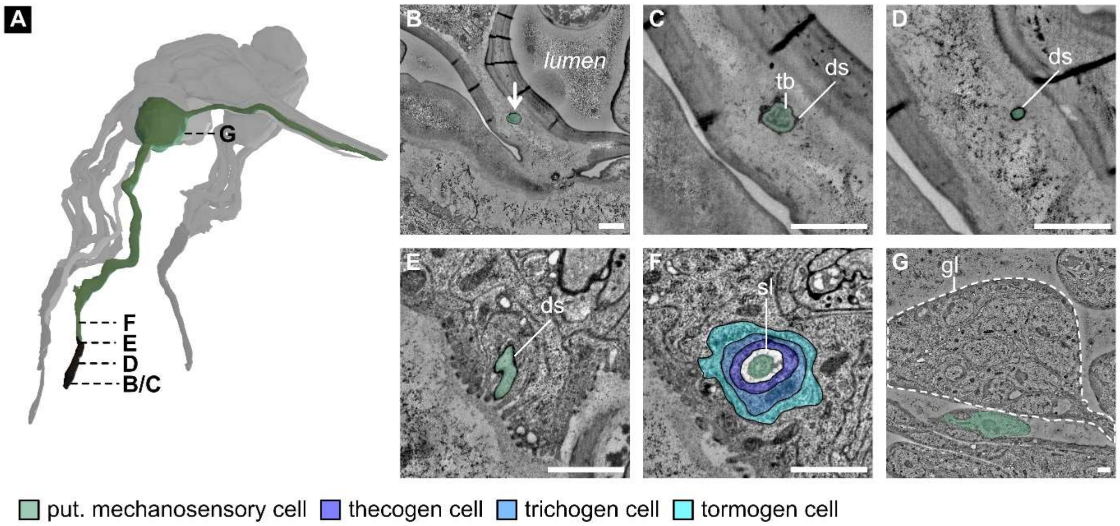


**Figure 3S1 – DPS-P/$S_1$. Internal morphology of the papilla/spot sensillum 1 (P/$S_1$) of the dorsal pharyngeal sensilla (DPS).**

(A) 3D reconstruction of the P/$S_1$ sensillum. A single sensory neuron (turquoise) innervates P/S1. (B-G) Serial STEM sections of the P/$S_1$ sensillum. (B) The P/S1 sensillum sits at the anterior end of the pharyngeal cavity (ph). (C) Its dendrite exhibits a tubular body at the tip. A dendritic sheath is present. The sensillum lacks a terminal pore. (D) Further proximal, the dendrite tapers down and thickens again (E). The dendrite is entirely enclosed by the dendritic sheath. (F) The dendrite is bathed in the sensillum lymph, but the dendritic sheath is absent at this level. The thecogen, trichogen and tormogen support cells are highlighted. (G) Section of the VPS ganglion. The cell body of the P/$S_1$ neuron is visible but lies beside the ganglion.

Abbreviations: ds - dendritic sheath; tb - tubular body; sl - sensillum lymph; gl - ganglion

Scale bars: B-G: 1 µm

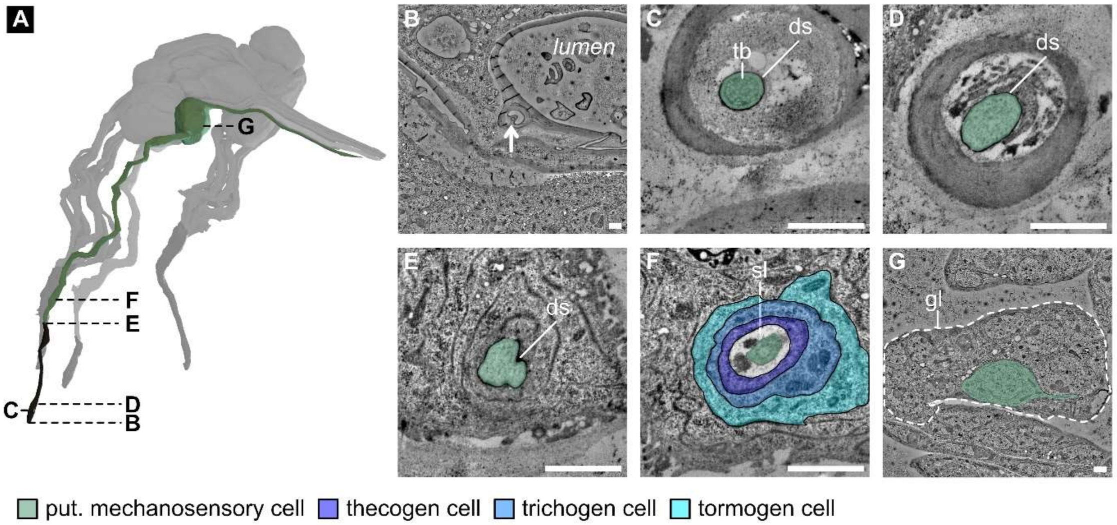


**Figure 3S2 – DPS-P/S2. Internal morphology of the papilla/spot sensillum 2 (P/$S_2$) of the dorsal pharyngeal sensilla (DPS).**

(A) 3D reconstruction of the P/$S_2$ sensillum. A single sensory neuron (turquoise) innervates S2. (B-G) Serial STEM sections of the P/$S_2$ sensillum. (B) The P/$S_2$ sensillum sits at the anterior end of the pharyngeal cavity (ph) in a deep "cuticle" channel (2,5 µm). (C) Its dendrite exhibits a tubular body at the tip. A dendritic sheath is present. The sensillum lacks a terminal pore. (D) Further proximal, the dendrite thickens (E). The dendritic sheath entirely encloses the dendrite. (F) The dendrite is bathed in the sensillum lymph, but the dendritic sheath is absent at this level. The thecogen, trichogen and tormogen support cells are highlighted. (G) Section of the VPS ganglion. The cell body of the P/$S_2$ neuron is visible and lies inside the ganglion.

Abbreviations: ds - dendritic sheath; tb - tubular body; sl - sensillum lymph; gl - ganglion

Scale bars: B-G: 1 µm

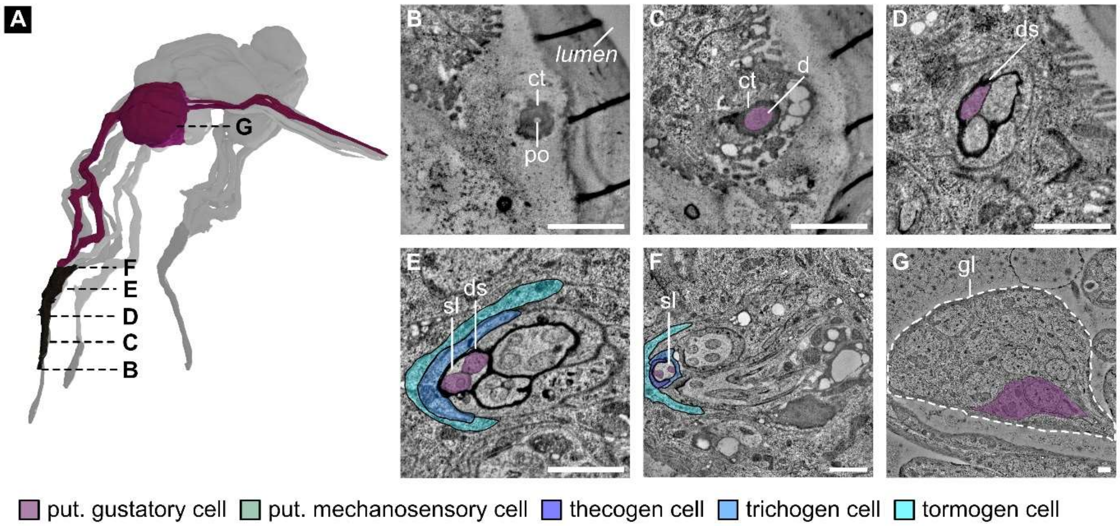


**Figure 3S3 – DPS-$T_1$. Internal morphology of the pit sensillum 1 ($T_1$) of the dorsal pharyngeal sensilla (DPS).**

(A) 3D reconstruction of the $T_1$ sensillum. Two sensory neurons (purple) innervate $T_1$. The sensillum shares a terminal pore to the pharyngeal lumen with pit sensilla $T_2$ and $T_3$ (see Figures 3S4 and 3S5). (B-G) Serial STEM sections of the $T_1$ sensillum. (B) The $T_1$ sensillum sits in a small pore at the anterior end of the pharyngeal cavity. (C) Its dendrites extend into the pore through the cuticle tube, together with the $T_2$ and $T_3$ dendrites. (D) Further proximal, the dendritic sheaths separate the dendrites of $T_1$ – $T_3$ from each other. (E) The dendrites' ciliary structures are visible and entirely enclosed by the dendritic sheath(s). The trichogen and tormogen support cells are visible. (F) The dendrites are bathed in the sensillum lymph, but the dendritic sheath disintegrates at this level. The thecogen, trichogen and tormogen support cells are visible. (G) Section of the VPS ganglion. The cell bodies of the $T_1$ neurons are visible and lie inside the ganglion.

Abbreviations: po - pore; ct - cuticle tube; d - dendrites; ds - dendritic sheath; sl - sensillum lymph; lm - lamellae; gl - ganglion

Scale bars: B-G: 1 µm

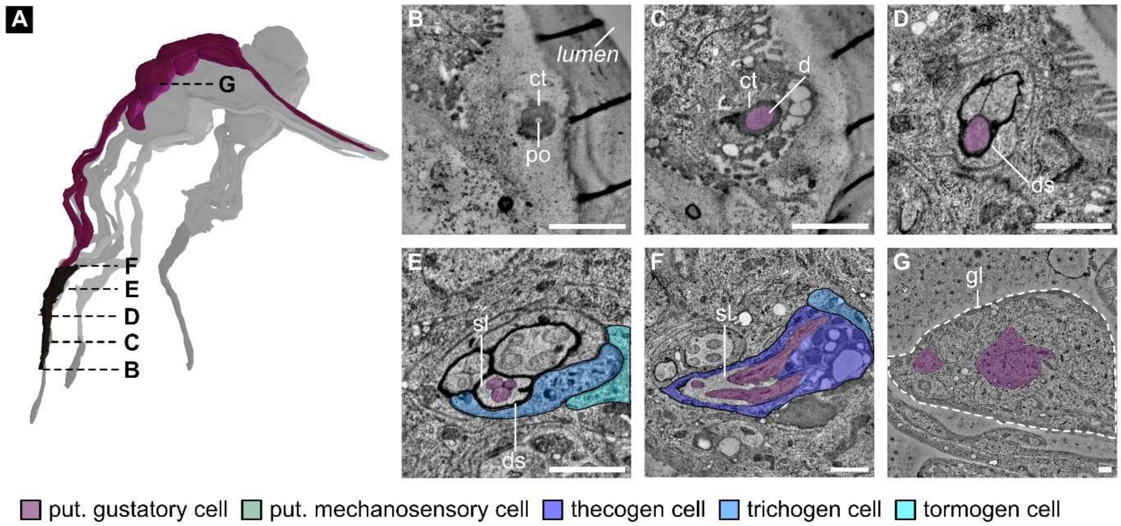


**Figure 3S4 – DPS-$T_2$. Internal morphology of the pit sensillum 2 ($T_2$) of the dorsal pharyngeal sensilla (DPS).**

(A) 3D reconstruction of the $T_2$ sensillum. Three sensory neurons (purple) innervate $T_2$. The sensillum shares a terminal pore to the pharyngeal lumen with pit sensilla $T_1$ and $T_3$ (see Figures 3S3 and 3S5). (B-G) Serial STEM sections of the $T_2$ sensillum. (B) The $T_2$ sensillum sits in a small pore at the anterior end of the pharyngeal cavity (ph). (C) Its dendrites extend into the pore through the cuticle channel, together with the $T_1$ and $T_3$ dendrites. (D) Further proximal, a dendritic sheath appears that separates the dendrites of $T_1$ – $T_3$ from each other. (E) The dendrites' ciliary structures are visible and entirely enclosed by the dendritic sheath(s). The trichogen and tormogen support cells are visible. (F) The dendrites are bathed in the sensillum lymph, but the dendritic sheath is entirely absent at this level. The thecogen and trichogen support cells are visible. (G) Section of the VPS ganglion. The cell bodies of the $T_2$ neurons are visible and lie inside the ganglion.

Abbreviations: po - pore; ct - cuticle tube; d - dendrites; ds - dendritic sheath; sl - sensillum lymph; lm - lamellae; gl - ganglion

Scale bars: B-G: 1 µm

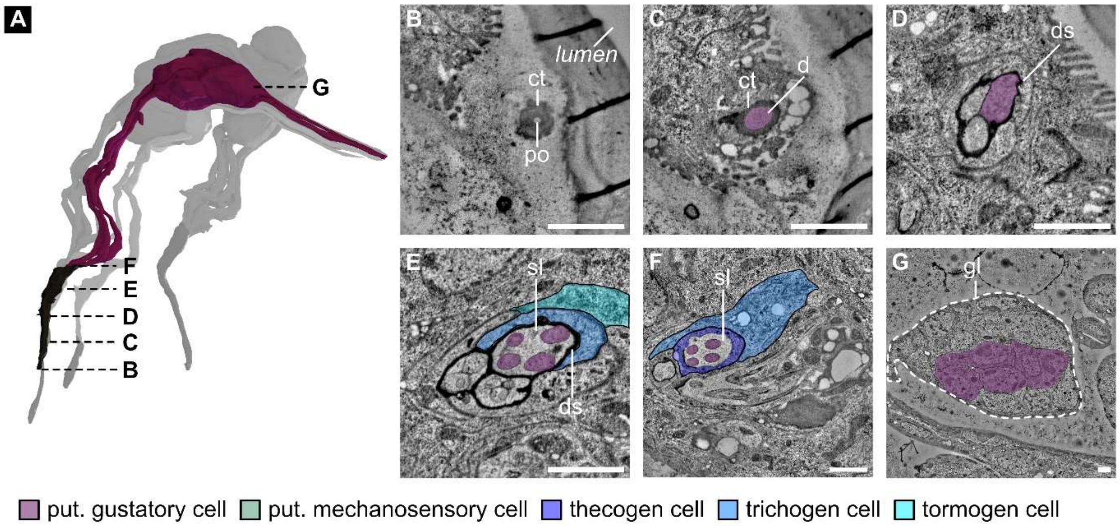


**Figure 3S5 – DPS-$T_3$. Internal morphology of the pit sensillum 3 ($T_3$) of the dorsal pharyngeal sensilla (DPS).**

(A) 3D reconstruction of the $T_3$ sensillum. Four sensory neurons (purple) innervate $T_3$. The sensillum shares a terminal pore to the pharyngeal lumen with pit sensilla $T_1$ and $T_2$ (see Figures 3S3 and 3S4). (B-G) Serial STEM sections of the $T_3$ sensillum. (B) The $T_3$ sensillum sits in a small pore at the anterior end of the pharyngeal cavity (ph). (C) Its dendrites extend into the pore through the cuticle channel, together with the $T_1$ and $T_2$ dendrites. (D) Further proximal, a dendritic sheath appears that separates the dendrites of $T_1$ - $T_3$ from each other. (E) The dendrites' ciliary structures are visible and entirely enclosed by the dendritic sheath(s). The trichogen and tormogen support cells are visible. (F) The dendrites are bathed in the sensillum lymph, but the dendritic sheath is entirely absent at this level. The thecogen and trichogen support cells are visible. (G) Section of the VPS ganglion. The cell bodies of the $T_3$ neurons are visible and lie inside the ganglion.

Abbreviations: po - pore; ct - cuticle tube; d - dendrites; ds - dendritic sheath; sl - sensillum lymph; lm - lamellae; gl - ganglion

Scale bars: B-G: 1 µm

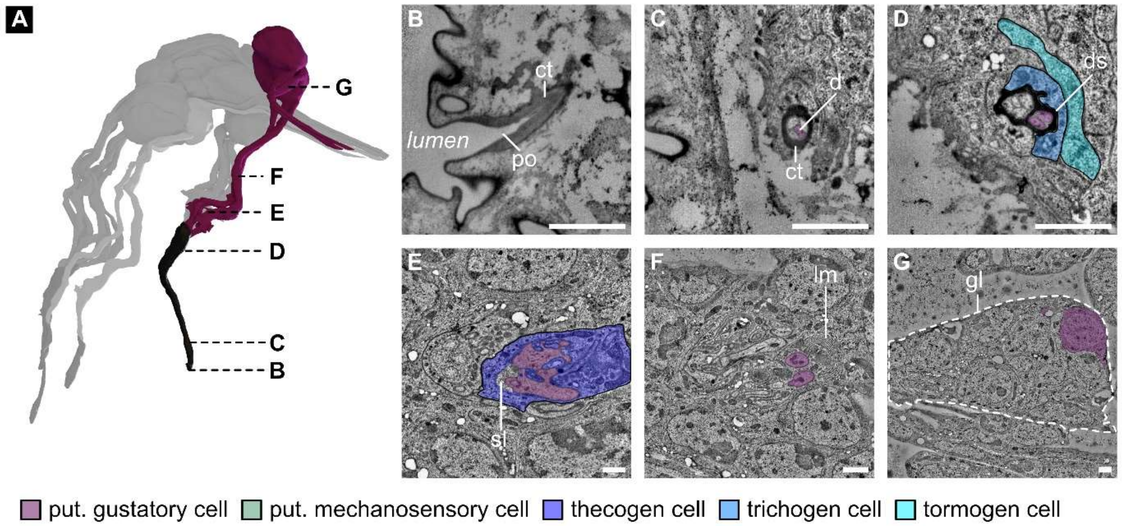


**Figure 3S6 – DPS-$T_4$. Internal morphology of the pit sensillum 4 ($T_4$) of the dorsal pharyngeal sensilla (DPS).**

(A) 3D reconstruction of the $T_4$ sensillum. Two sensory neurons (purple) innervate $T_4$. The sensillum shares a terminal pore to the pharyngeal lumen with the pit sensillum $T_5$ (see Figure 3S7). (B-G) Serial STEM sections of the $T_4$ sensillum. (B) The $T_4$ sensillum sits in a small pore at the posterior end of the pharyngeal cavity (ph). (C) Its dendrites extend into the pore through the cuticle channel, together with the $T_5$ dendrites. (D) Further proximal, a dendritic sheath appears that separates the dendrites of $T_4$ and $T_5$ from each other. The trichogen and tormogen support cells are visible. (E) The dendrites ciliary structures and the thecogen cell are visible; the dendritic sheaths are absent at this level. The dendrites are bathed in the sensillum lymph. (F) The sheath cell is highly lamellated at this level. (G) Section of the VPS ganglion. One cell body and one dendrite of the $T_4$ neurons are visible and lie inside the ganglion.

Abbreviations: po - pore; ct - cuticle tube; d - dendrites; ds - dendritic sheath; sl - sensillum lymph; lm - lamellae; gl - ganglion

Scale bars: B-G: 1 µm

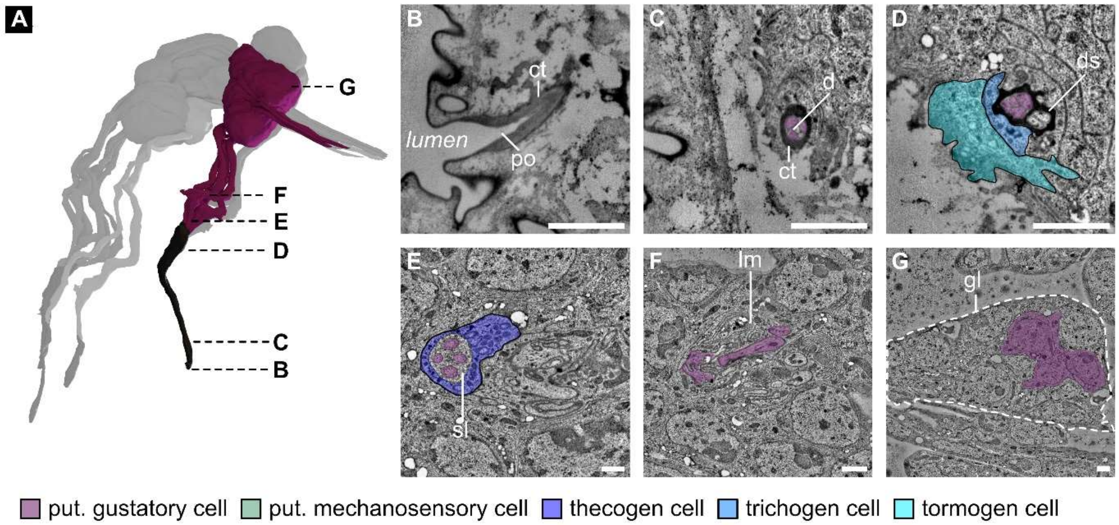


**Figure 3S7 – DPS-$T_5$. Internal morphology of the pit sensillum 5 ($T_5$) of the dorsal pharyngeal sensilla (DPS).**

(A) 3D reconstruction of the $T_5$ sensillum. Four sensory neurons (purple) innervate $T_5$. The sensillum shares a terminal pore to the pharyngeal lumen with the pit sensillum $T_4$ (see Figure 3S6). (B-G) Serial STEM sections of the $T_5$ sensillum. (B) The $T_5$ sensillum sits in a small pore at the posterior end of the pharyngeal cavity (ph). (C) Its dendrites extend into the pore through the cuticle channel, together with the $T_4$ dendrites. (D) Further proximal, a dendritic sheath appears that separates the dendrites of $T_4$ and $T_5$ from each other. The trichogen and tormogen support cells are visible. (E) The dendrites ciliary structures and the thecogen cell are visible; the dendritic sheath is absent at this level. The dendrites are bathed in the sensillum lymph. (F) The sheath cell is highly lamellated at this level. (G) Section of the VPS ganglion. Three cell bodies and one dendrite of the $T_5$ neurons are visible and lie inside the ganglion.

Abbreviations: po - pore; ct - cuticle tube; d - dendrites; ds - dendritic sheath; sl - sensillum lymph; lm - lamellae; gl - ganglion

Scale bars: B-G: 1 µm

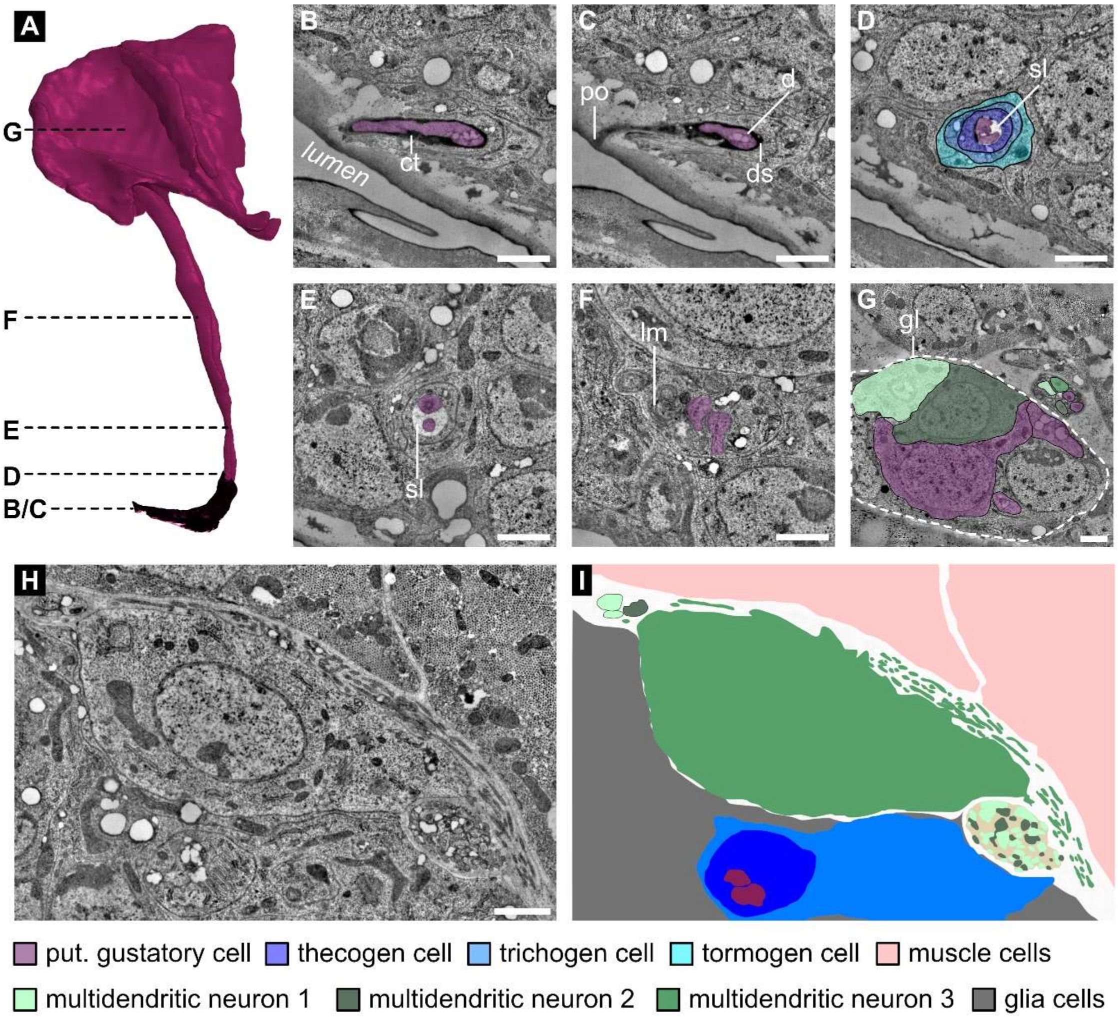


**Figure 4S1 – DPO-T1. Internal morphology of the pit sensillum 1 ($T_1$) of the dorsal pharyngeal organ (DPO).**

(A) 3D reconstruction of the $T_1$ sensillum. Two sensory neurons (purple) innervate $T_1$. (B-G) Serial STEM sections of the $T_1$ sensillum. (B) The terminal pore is visible. (C) The pore is connected to a cuticle channel, which is innervated by the dendrites (D). The two ciliary structures (purple) are visible. A dendritic sheath is present but disintegrating at this level. The tips of the dendrites are bathed in the sensillum lymph, enclosed by the thecogen, trichogen, and tormogen support cells. (E) The two ciliary structures (purple) are visible. The dendritic sheath is absent at this level. (F) The outer dendritic segments pass through the outer support cells. (G) Section of the DPO ganglion (DPOG). The cell bodies of the $T_1$ neurons and of multidendritic neurons A and B are visible. (H+I) Section of the DPO ganglion (DPOG); the cell body of multidendritic neuron C is visible, from which multiple dendrites extend in the extracellular space towards the muscle cells.

Abbreviations: po - pore; ct - cuticle tube; d - dendrites; ds - dendritic sheath; sl - sensillum lymph; lm - lamellae; gl - ganglion

Scale bars: B-H: 1 µm

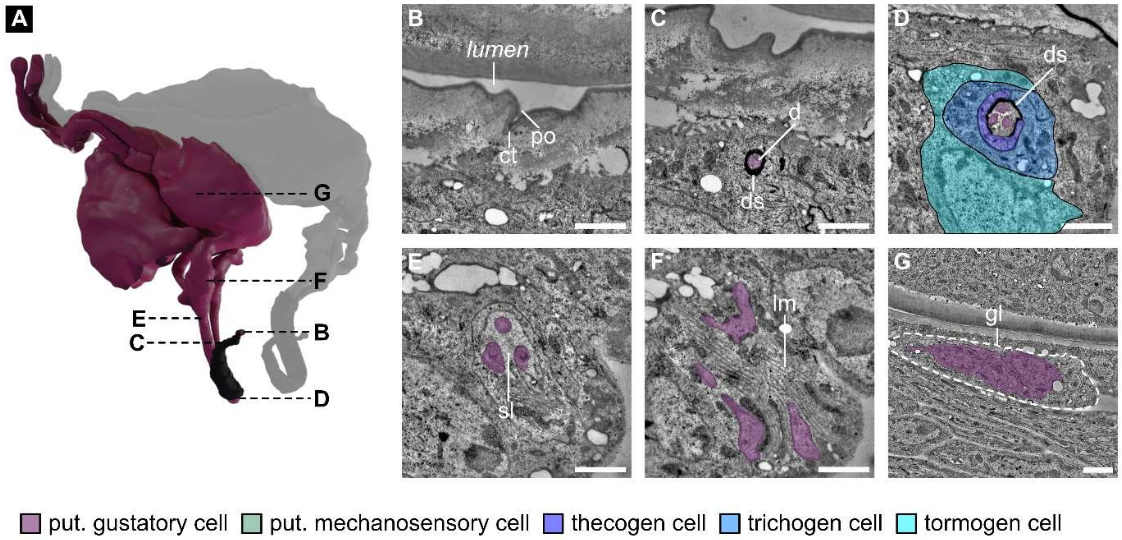


**Figure 4S2 – PPS-T$_1$. Internal morphology of the pit sensillum 1 (T$_1$) of the posterior pharyngeal organ (PPS).**

(A) 3D reconstruction of the T$_1$ sensillum. Three sensory neurons (purple) innervate T$_1$. (B-G) Serial STEM sections of the T$_1$ sensillum. (B) The terminal pore is visible. (C) The pore is connected to a cuticle channel, which is innervated by the dendrites (D). A dendritic sheath encloses the dendrites but disintegrates at this level. The tips of the dendrites are bathed in the sensillum lymph, enclosed by the thecogen, trichogen and tormogen support cells. (E) The three ciliary structures (purple) are visible. The dendritic sheath is entirely absent at this level. (F) The outer dendritic segments pass through the thecogen cell, which is highly lamellated. (G) Section of the PPS ganglion (PPSG). The cell bodies of the T$_1$ neurons are visible.

Abbreviations: po - pore; ct - cuticle tube; d - dendrites; ds - dendritic sheath; sl - sensillum lymph; lm - lamellae; gl - ganglion

Scale bars: B-G: 1 µm

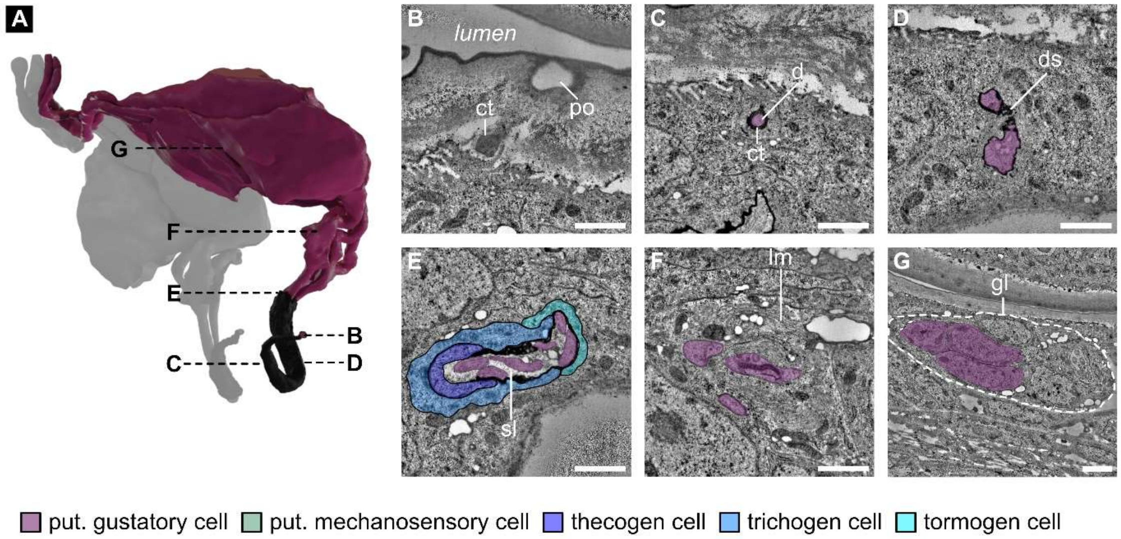


**Figure 4S3 – PPS-$T_2$. Internal morphology of the pit sensillum 2 ($T_2$) of the posterior pharyngeal organ (PPS).**

(A) 3D reconstruction of the $T_2$ sensillum. Three sensory neurons (purple) innervate $T_2$. (B-G) Serial STEM sections of the $T_2$ sensillum. (B) The terminal pore is visible. (C) The pore is connected to a cuticle channel, which is innervated by the dendrites (D). The dendrites are enclosed by a dendritic sheath (E). The three ciliary structures (purple) are visible. A dendritic sheath is present but disintegrating at this level. The tips of the dendrites are bathed in the sensillum lymph and are enclosed by the thecogen, trichogen, and tormogen support cells. (F) The outer dendritic segments pass through the thecogen cell, which is highly lamellated. (G) Section of the PPS ganglion (PPSG). The cell bodies of the $T_2$ neurons are visible.

Abbreviations: po - pore; ct - cuticle tube; d - dendrites; ds - dendritic sheath; sl - sensillum lymph; lm - lamellae; gl - ganglion

Scale bars: B-G: 1 µm

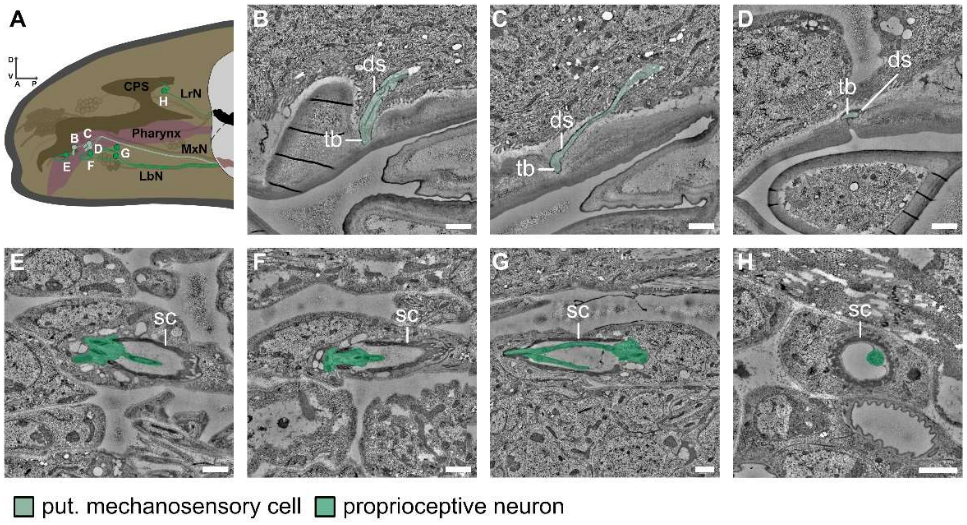


**Figure 4S4 – other. Internal morphology of the papilla/spot sensilla and chordotonal organs associated with the feeding apparatus.**

(A) Schematic drawing of the larval head region highlighting the position of the additional papilla/spot sensilla and chordotonal organs.

(B) – (D) EM images of the dendritic parts of the three papilla/spot sensilla inserting into the pharyngeal lumen.

(E) – (F) EM images of the dendritic parts of the four chordotonal organs associated with the feeding apparatus.

Sensilla were identified using prominent structures like a dendritic sheath and a tubular body (papilla/spot sensilla) or a sensory neuron inserted into a scolopale (chordotonal organs.

Scale bars: B-H: 1 µm

Abbreviations: D - dorsal; V - ventral; A - anterior; P - posterior; CPS - cephalopharyngeal skeleton; LrN – labral nerve; MxN – maxillary nerve; LbN – labial nerve; ds - dendritic sheath; tb - tubular body; sc - scolopale
